# “Is Voice A Marker for Autism Spectrum Disorder? A Systematic Review And Meta-Analysis”

**DOI:** 10.1101/046565

**Authors:** Riccardo Fusaroli, Anna Lambrechts, Dan Bang, Dermot M Bowler, Sebastian B Gaigg

**Author notes:** Corresponding author: Riccardo Fusaroli, address: Jens Christian Skous vej 2, 8000 Aarhus Denmark. Grant sponsor: Interacting Minds Center; Grant ID: Clinical Voices. The authors declare that no conflict of interest exists.

## Abstract

**Lay Abstract:** Individuals with Autism Spectrum Disorder (ASD) are reported to speak in distinctive ways. Distinctive vocal production should be better understood as it can affect social interactions and social development and could represent a noninvasive marker for ASD. We systematically review the existing scientific literature reporting quantitative acoustic analysis of vocal production in ASD and identify repeated and consistent findings of higher pitch mean and variability but not of other differences in acoustic features. We also identify a recent approach relying on multiple aspects of vocal production and machine learning algorithms to automatically identify ASD from voice only. This latter approach is very promising, but requires more systematic replication and comparison across languages and contexts. We outline three recommendations to further develop the field: open data, open methods, and theory-driven research.

**Scientific Abstract:** Individuals with Autism Spectrum Disorder (ASD) tend to show distinctive, atypical acoustic patterns of speech. These behaviours affect social interactions and social development and could represent a non-invasive marker for ASD. We systematically reviewed the literature quantifying acoustic patterns in ASD. Search terms were: (prosody OR intonation OR inflection OR intensity OR pitch OR fundamental frequency OR speech rate OR voice quality OR acoustic) AND (autis* OR Asperger). Results were filtered to include only: empirical studies quantifying acoustic features of vocal production in ASD, with a sample size > 2, and the inclusion of a neurotypical comparison group and/or correlations between acoustic measures and severity of clinical features. We identified 34 articles, including 30 univariate studies and 15 multivariate machine-learning studies. We performed meta-analyses of the univariate studies, identifying significant differences in mean pitch and pitch range between individuals with ASD and comparison participants (Cohen's d of 0.4–0.5 and discriminatory accuracy of about 61–64%). The multivariate studies reported higher accuracies than the univariate studies (63–96%). However, the methods used and the acoustic features investigated were too diverse for performing meta-analysis. We conclude that multivariate studies of acoustic patterns are a promising but yet unsystematic avenue for establishing ASD markers. We outline three recommendations for future studies: open data, open methods, and theory-driven research.

## 1. Introduction

From its earliest characterizations, ASD has been associated with peculiar tones of voice and disturbances of prosody (Asperger, 1944; Goldfarb, Braunstein, & Lorge, 1956; Kanner, 1943; Pronovost, Wakstein, & Wakstein, 1966;Simmons & Baltaxe, 1975). Although 70–80% of individuals with ASD develop functional spoken language, at least half of the ASD population displays early atypical acoustic patterns (Paul et al., 2005a; Rogers et al., 2006; Shriberg et al., 2001), which persist while other aspects of language improve (Baltaxe & Simmons, 1985; Depape, Chen, Hall, & Trainor, 2012). These atypical acoustic patterns have been qualitatively described as flat, monotonous, variable, sing-songy, pedantic, robot-or machine-like, hollow, stilted or exaggerated and inappropriate (Amorosa, 1992; Baltaxe, 1981; Depape, et al., 2012; Järvinen-Pasley, Peppé, King-Smith, & Heaton, 2008; Lord, Rutter, & Le Couteur, 1994). Such distinctive vocal characteristics are one of the earliest-appearing markers of a possible ASD diagnosis (Oller et al., 2010; Paul, Fuerst, Ramsay, Chawarska, & Klin, 2011; Warlaumont, Richards, Gilkerson, & Oller, 2014).

An understanding of vocal production in ASD is important because acoustic abnormalities may play a role in the social-communicative impairments associated with the disorder (Depape, et al., 2012; Klopfenstein, 2009). For example, individuals with ASD have difficulties with the communication of affect (Travis & Sigman, 1998) - which relies on the production of prosodic cues - leading to negative social judgments on the part of others (Fay & Schuler, 1980; Paul et al., 2005b; Shriberg, et al., 2001; Van Bourgondien & Woods, 1992) and in turn social withdrawal and social anxiety (Alden & Taylor, 2004). Such disruption of communication and interaction may have long-term effects, compromising the development of social-communicative abilities (Warlaumont, et al., 2014).

Atypical prosody is already considered a marker for ASD in gold-standard diagnostic assessments such as the Autism Diagnostic Observation Schedule (Lord, et al., 1994), and recent evidence indicates that speech in ASD may be characterized by relatively unique acoustic features that can be quantified objectively (Bone et al., 2013; Fusaroli, Lambrechts, Yarrow, Maras, & Gaigg, 2015; Oller, et al., 2010). Prosody production has also been argued to be a “bellwether” behavior that can serve as a marker of the specific cognitive and social functioning profile of an individual (Bone et al., 2014; Diehl, Berkovits, & Harrison, 2010; Paul, et al., 2005a). Such diagnostic profiling is especially needed now that the diagnosis of ASD (since the publication of the DSM-5) pools together previously distinct disorders (e.g., Asperger syndrome and childhood disintegrative disorder).

Studies of prosody in ASD can be grouped according to four key aspects of speech production: pitch, volume, duration and voice quality (Cummins et al., 2015; Titze, 1994). The speech of individuals with ASD has been described as monotone, as having inappropriate pitch and pitch variation (Baltaxe, 1984; Fay & Schuler, 1980; Goldfarb, Goldfarb, Braunstein, & Scholl, 1972; Paccia & Curcio, 1982; Pronovost, et al., 1966) and as being too loud or too quiet, sometimes inappropriately shifting between the two (Goldfarb, et al., 1972; Pronovost, et al., 1966; Shriberg, Paul, Black, & van Santen, 2011; Shriberg, et al., 2001). Further, individuals with ASD have been reported to speak too quickly or too slowly (Baltaxe, 1981; Goldfarb, et al., 1972; Simmons & Baltaxe, 1975) and many descriptions of their speech have highlighted a distinctive voice quality characterized as “hoarse”, “harsh” and “hyper-nasal” (Baltaxe, 1981; Pronovost, et al., 1966), with a higher recurrence of squeals, growls, and yells (Sheinkopf, Mundy, Oller, & Steffens, 2000).

The research evidence is diverse, in terms of both methods and interpretations. An early review of 16 qualitative studies of speech in ASD found it difficult to draw any firm conclusions (McCann & Peppé, 2003). Shortcomings of the reviewed studies were: (1) small sample size; (2) underspecified criteria for the (qualitative) descriptions of speech production; (3) lack of quantitative measures of speech production; (4) use of heterogeneous and non-standardized tasks; and (5) little theory-driven research. Since that review, the literature on prosody in ASD has grown substantially, particularly with respect to the use of signal-processing techniques that overcome some of the limitations involved in qualitative studies (Banse & Scherer, 1996; Grossman, Bemis, Skwerer, & Tager-Flusberg, 2010). The purpose of the present paper is to provide a systematic and critical review of recent research on the acoustic quantitative characteristics of speech production in ASD. This focus ensures minimal overlap with the literature reviewed by McCann & Peppé (2003) and is motivated by the more general question of whether automated speech-processing procedures can be used in the diagnosis of ASD.

We identified two different groups of studies: univariate studies and multivariate machine-learning studies. Univariate studies seek to identify differences between ASD and comparison groups by investigating one acoustic feature at a time. In contrast, multivariate machine-learning studies use multiple features (multivariate) to build statistical models that can classify previously unheard voice samples into ASD and comparison groups (machine-learning).

A particular focus of this review will be whether acoustic characteristics of speech production can be used as markers of ASD, that is, as a directly measurable index derived from sensitive and reliable quantitative procedures that is associated with the condition and/or its clinical features (e.g. Ruggeri et al, 2014). Since ASD involves a high degree of heterogeneity of clinical features and their severity, it is crucial to assess how widely acoustic markers can apply to a wide range of individuals with ASD, and whether the markers reflect severity and progression of clinical features over time (e.g. in the context of intervention programs or aging). It should also be emphasized that, in light of the heterogeneity of individuals with ASD and the need for a reliable marker of ASD, the review will not speculate on the significance of the findings of isolated studies. Instead, the focus will be on finding patterns across studies, which are more likely to generalize to new samples (Yarkoni & Westfall, 2016).

The review will be structured as follows. Section 2 will define the search and selection criteria for the literature review. Sections 3 and 4 will present the results of the review. Section 3 focuses on univariate studies and, where more than five studies focused on the same feature, provides meta-analyses of the effect sizes. Section 4 focuses on multivariate studies and in particular the attempt to use machine-learning techniques to develop acoustic markers of ASD. We end by critically assessing the findings and advancing recommendations for future research.

## 2. Methods: The criteria for the literature search

A literature search was conducted using Google Scholar, PubMed and Web of Science on April 15 2015, updated on March 4 2016 and then again on June 21 2016. The search terms used were (prosody OR intonation OR inflection OR intensity OR pitch OR fundamental frequency OR speech rate OR voice quality OR acoustic) AND (autis^*^ OR Asperger). Additional search for unpublished studies was performed through additional web searches (on Google and Bing), and by directly contacting authors of the published studies and interested participants of the IMFAR 2014, 2015 and 2016 conferences. Furthermore it should be noted that Google Scholar covers most (if not all) dissertation repositories. The papers thus found were searched for additional references and the resulting set was screened by two of the authors (RF and AL) according to the following criteria: empirical study, quantification of acoustic features in the vocal production of participants with ASD, sample including at least two individuals with ASD, inclusion of a typically developing comparison group (TD) or an assessment of variation in acoustic features in relation to severity of clinical features. Non-TD comparison groups (e.g. with language impairment, or ADHD) were not included as not enough studies were present to assess patterns beyond the single study.

For all resulting papers we report sample sizes for ASD and TD groups, matching criteria, age, verbal and non-verbal level of function, speech production task, results and estimates of the acoustic measures (mean and standard deviation) if available, in dedicated tables (see Tables 1 to 5). To facilitate comparison between studies, the vocal production tasks were grouped into three categories. The first category, *constrained production,* includes tasks such as reading aloud and repeating linguistic stimuli. In this category, the focus is on the form of speech production, more than on its contents (e.g. the actual words and meaning expressed). The second category, *spontaneous production*, includes tasks such as free description of pictures and videos or telling stories. This category of tasks involves a more specific focus on the contents of speech production. The third category, *social interaction,* includes spontaneous and semi-structured conversations such as ADOS interviews. This category adds a stronger emphasis on social factors and interpersonal dynamics.

We extracted statistical estimates (mean and standard deviation for the ASD and TD groups) of the features when available and contacted the corresponding authors of the articles that did not provide these statistics^1^. When this process yielded statistical estimates of one feature from at least five independent studies, we ran a meta-analysis to estimate an overall effect size - that is, a weighted standardized mean difference (Cohen's d) between the ASD and the TD groups for univariate studies and sensitivity/specificity of classification for the multivariate machine-learning studies. We note that only the univariate studies provided enough data to perform meta-analyses.

Meta-analyses were performed following well-established procedures detailed in (Doebler & Holling, 2015; Field & Gillett, 2010; Quintana, 2015; Viechtbauer, 2010). We first calculated the size (Cohen's d), statistical significance (p-value) and overall variance (or τ^2^) of effects observed across studies. We then assessed whether the overall variance could be explained by within-study variance (e.g., due to measurement noise or heterogeneity in the ASD samples included in the studies) using Cochran's Q (Cochran, 1954) and I^2^ statistics (Higgins, Thompson, Deeks, & Altman, 2003). Third, we assessed whether systematic factors - speech production task (constrained production, spontaneous production, social interaction) and language employed in the task (e.g. American English, or Japanese) - could further explain the overall variance. Age would be a third crucial factor to add to the analysis. However, the studies analyzed spanned wide age ranges, which did not allow making any clear division in age groups (such as childhood, adolescence and adulthood). Finally, we investigated the effect of influential studies (single studies strongly driving the overall results) and publication bias (tendency to write up and publish only significant findings, ignoring null findings and making the literature unrepresentative of the actual population studied) on the robustness of our analysis. This was estimated using rank correlation tests assessing whether lower sample sizes (and relatedly higher standard error) were related to bigger effect sizes. A significant rank correlation indicates a likely publication bias and inflated effect sizes due to small samples. All analyses were performed using the metafor v.1.9.8 and mada v.0.5.7 packages in R 3.3. All data and R-code employed are available at https://github.com/fusaroli/AcousticPatternsInASD and on FigShare with the doi: https://dx.doi.org/10.6084/m9.figshare.3457751.v2 (Fusaroli, 2016).

## 3. Results

### 3.1. Literature search results

The initial literature screening yielded 108 papers discussing prosody and voice in ASD. The second stricter screening yielded 34 papers, with each paper sometimes reporting more than one study. In total, our primary literature included 30 univariate studies and 15 multivariate machine-learning studies. The remaining 74 papers (qualitative studies, theory or reviews) were used as background literature only and cited when relevant.

### 3.2. Differences in acoustic patterns between ASD and comparison populations (univariate studies)

#### 3.2.1 Pitch

Pitch reflects the frequency of vibrations of the vocal cords during vocal production. During vocal production, individuals often modulate their pitch to convey pragmatic or contextual meaning: for example, marking an utterance as having an imperative, declarative or ironic intent, or even to express emotions (Banse & Scherer, 1996; Bryant, 2010; Fusaroli & Tylén, 2016; Michael et al., 2015; Mushin, Stirling, Fletcher, & Wales, 2003).

Our literature screening yielded 24 studies employing acoustic measures of pitch (see Tables 1–2). Five summary statistics were used: mean, standard deviation (SD), range (defined between highest and lowest pitch), mean absolute deviation from the median (a measure of variability especially robust to outliers) and coefficient of variation (standard deviation divided by mean). Some researchers also quantified the temporal trajectory or profile of pitch, estimating the slope (ascending, descending or flat) of pitch over time (Bone, et al., 2014; Green & Tobin, 2009). We report the latter measures when the signal-processing is automated and does not rely on manual coding.

**Table 1.**
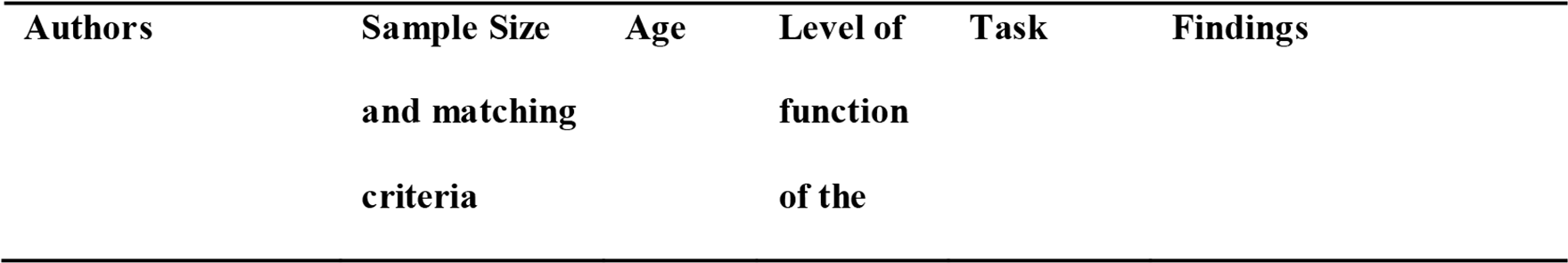
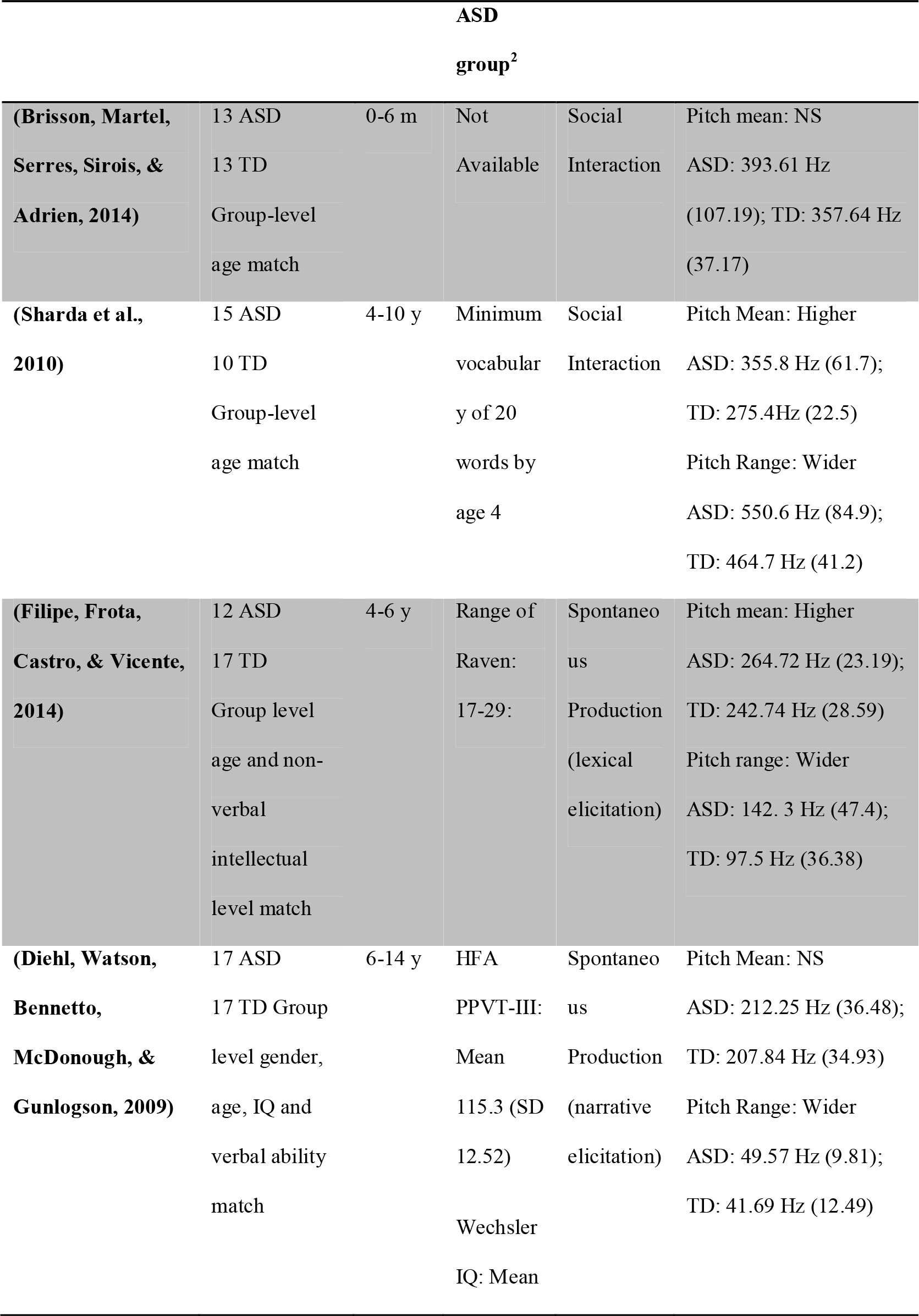
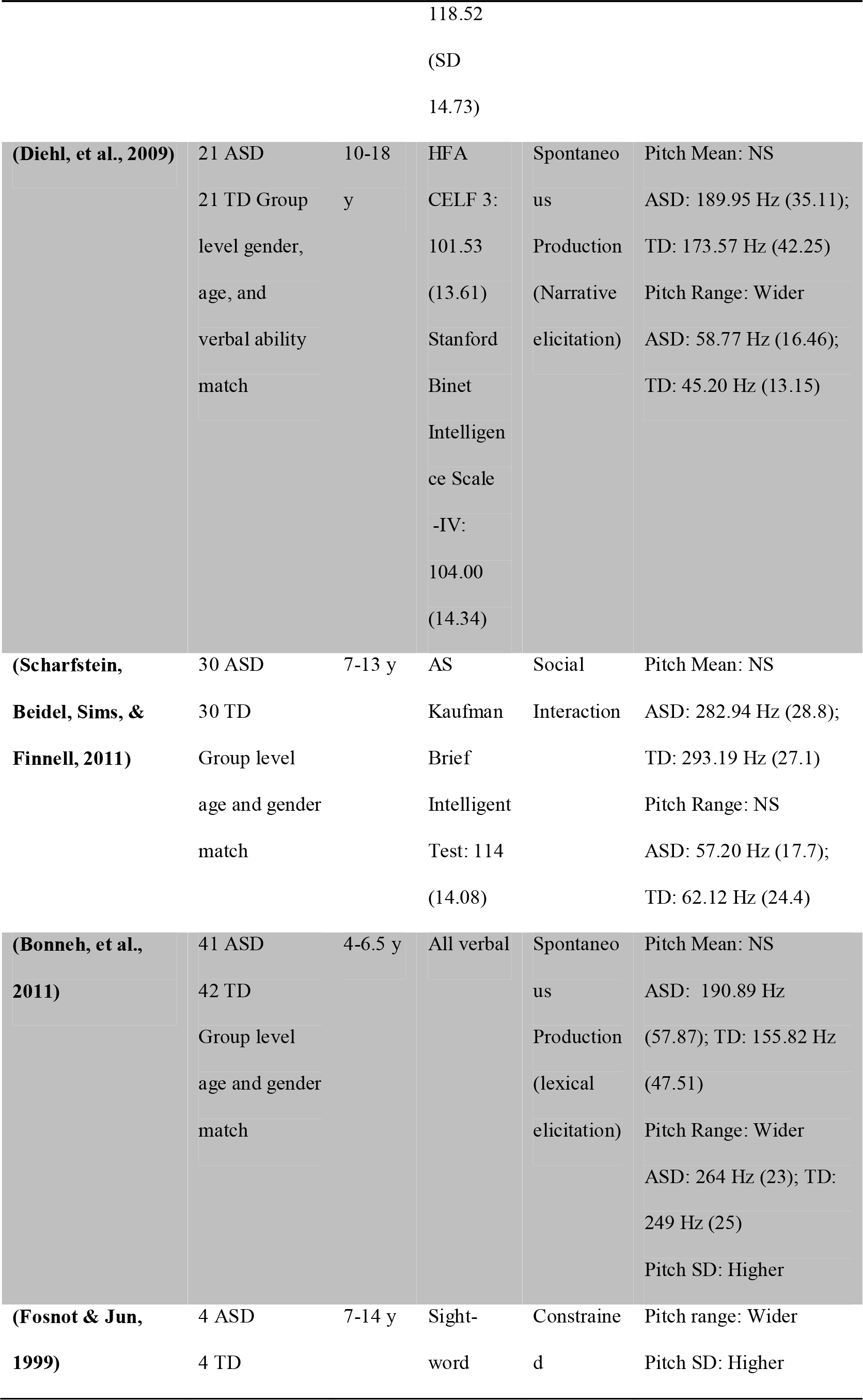
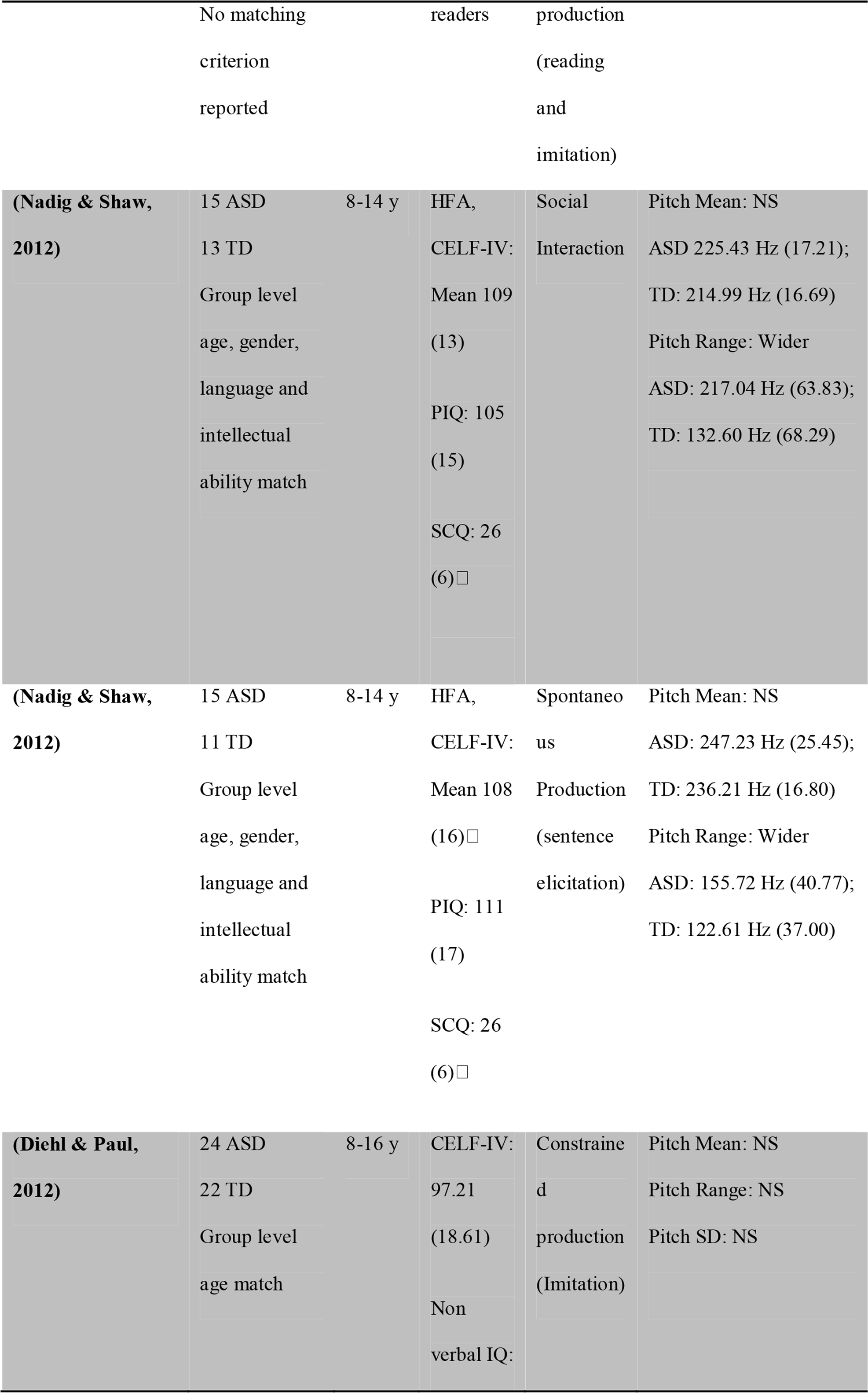
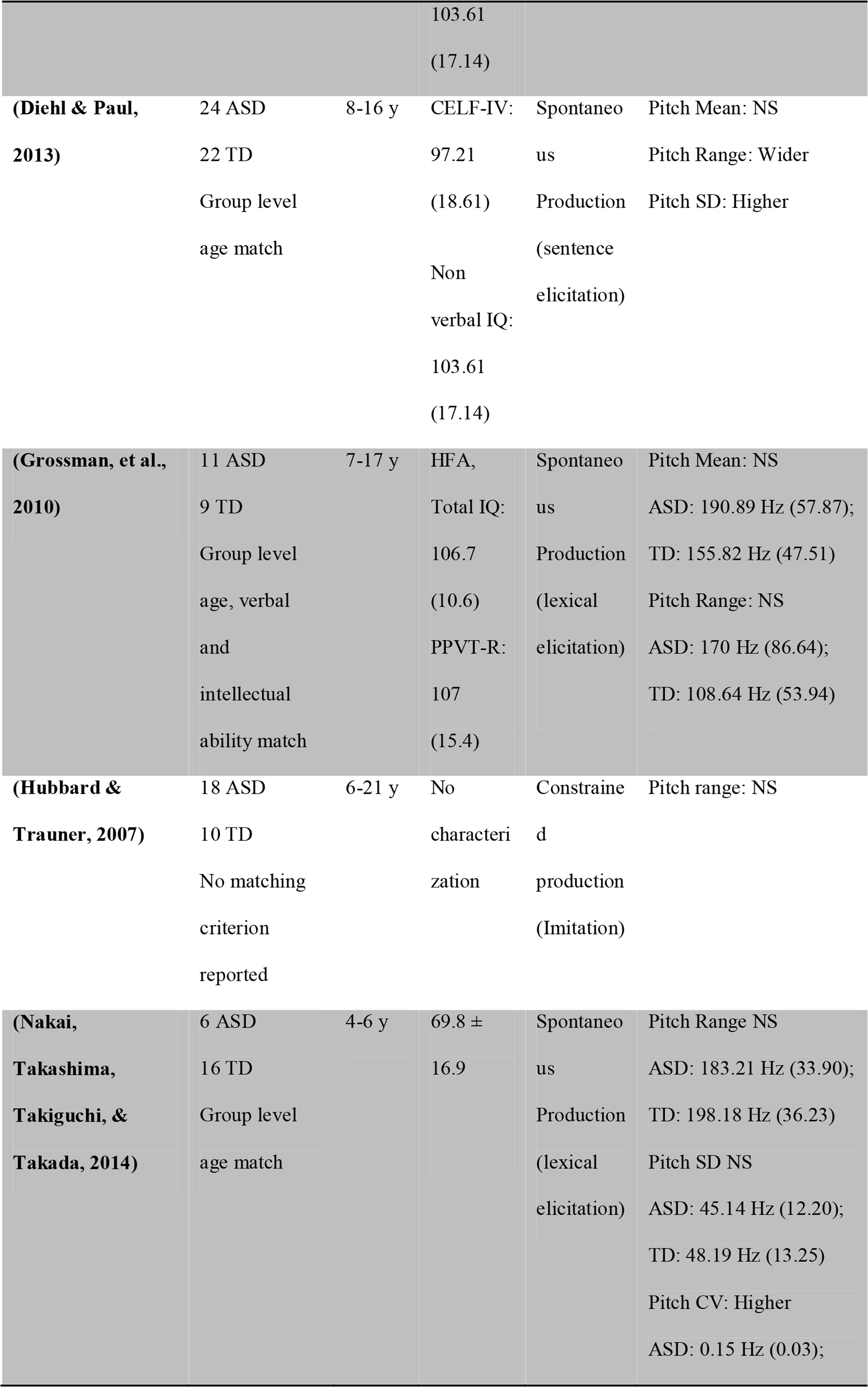
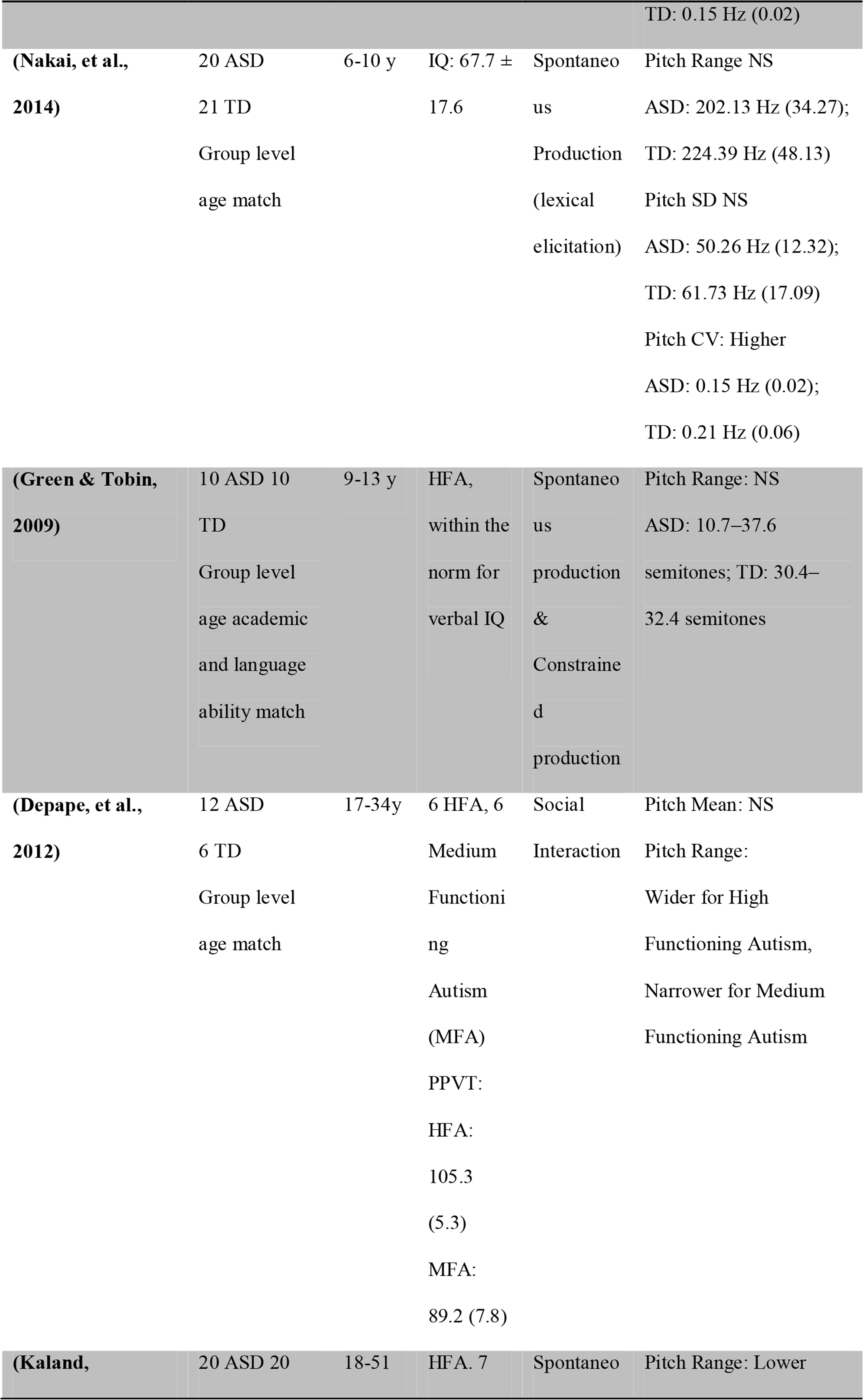
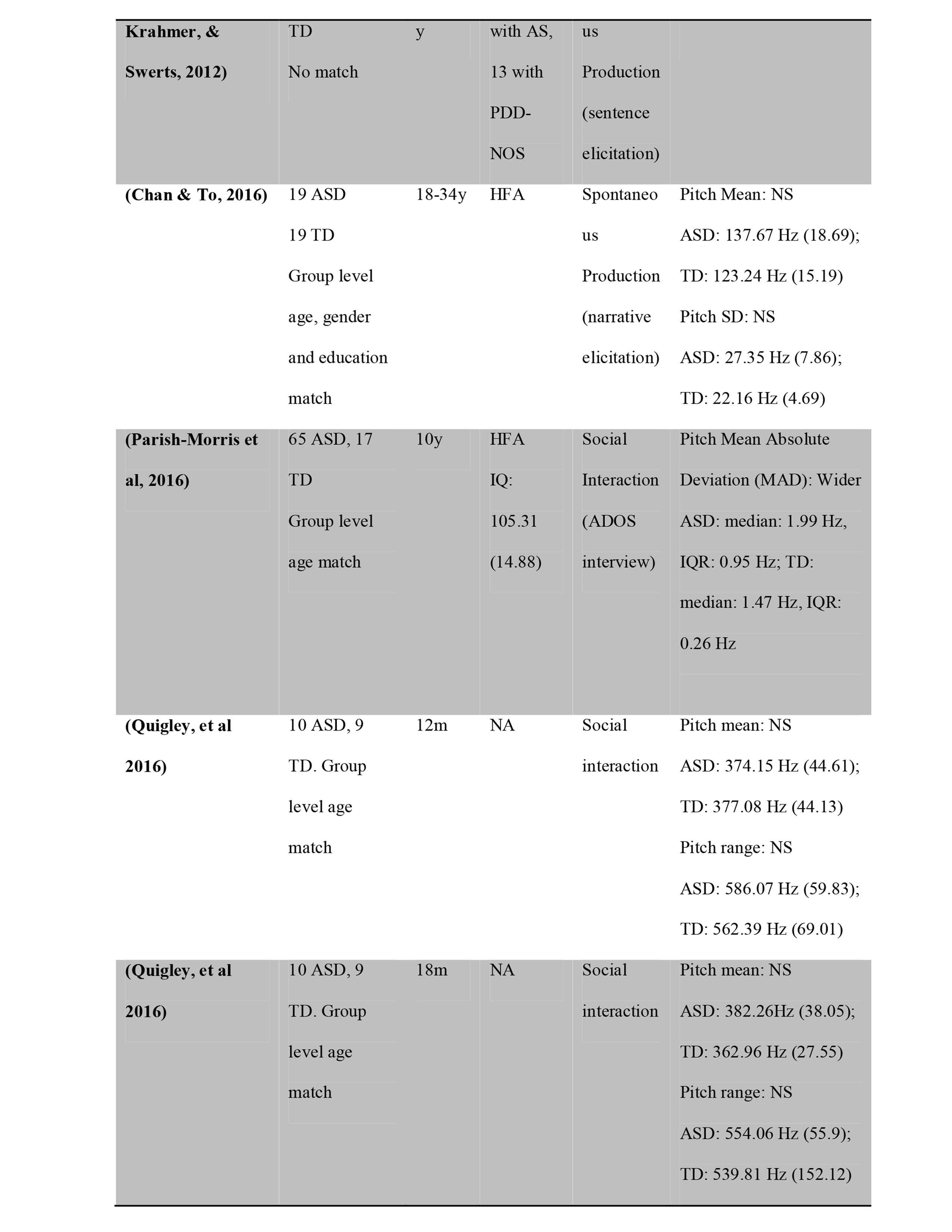
*Summary statistics of the pitch properties of ASD and TD groups in each study. When present, or provided by the authors, mean and standard deviation (in parenthesis) of the summary statistics are reported. NS: Non-significant difference between groups*^2^.

*Pitch mean* was investigated in 16 studies (255 participants with ASD and 239 comparison participants). Only two of these studies reported a significant group difference with higher pitch mean in the ASD groups (Filipe, et al., 2014; Sharda, et al., 2010). The remaining 14 studies report null findings. The meta-analysis included 11 studies for a total of 219 participants with ASD and 211 comparison participants (see Figure 1). The overall estimated difference (Cohen's d) in mean pitch between the ASD and TD groups was 0.41 (95% CIs: 0.15 0.68, p=0.003) with an overall variance (τ^2^) of 0.1 (95% CIs: 0 0.48). Much of the variance (I^2^: 44.11%, 95% CIs: 0 79.53) could not be reduced to random sample variability between studies (Q-stats=21.35, p=0.046). However, neither task (estimate: 0.09, 95% CIs −0.46 0.63, p=0.76) nor language (estimate: 0.05, 95% CIs −0.04 0.13, p=0.26) could significantly explain it.

**Figure 1.**
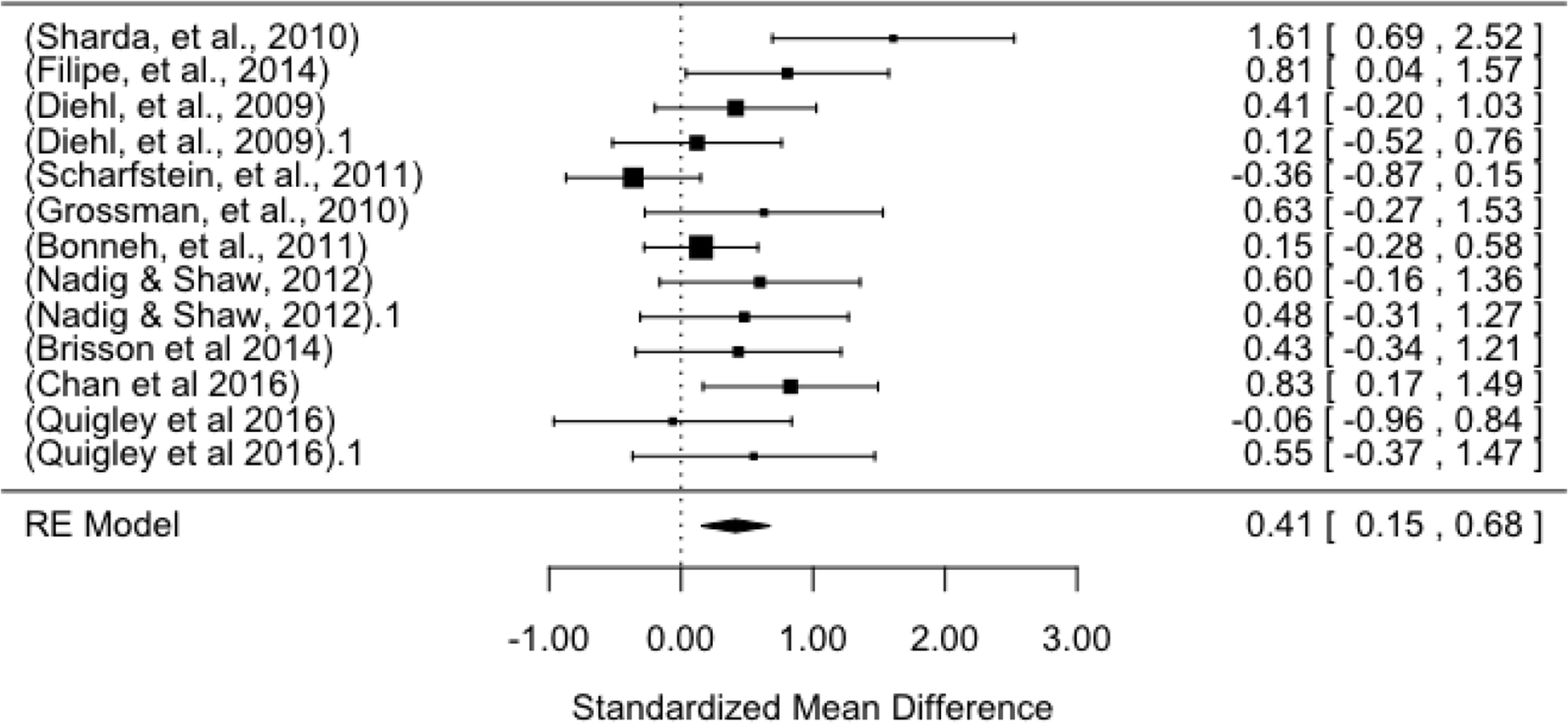
Forest plot of effect sizes (Cohen's d) in pitch mean between the ASD and comparison populations. The x-axis reports the effect size (positive values indicate higher mean pitch in ASD, while negative lower) and the y-axis the studies for which statistical estimates of pitch mean were provided. The dotted vertical line indicates the null hypothesis (no difference between the populations).

One study (Sharda, et al., 2010) with a large effect size and large standard error significantly drives the overall effect (see the lowest right point in Figure 2). Removing this study yielded a smaller but still significant overall effect size (0.33, 95% CIs 0.09 0.56, p=0.006). The data did not reveal any likely publication bias (Kendall's τ=0.36, p=0.1; Figure 2).

**Figure 2.**
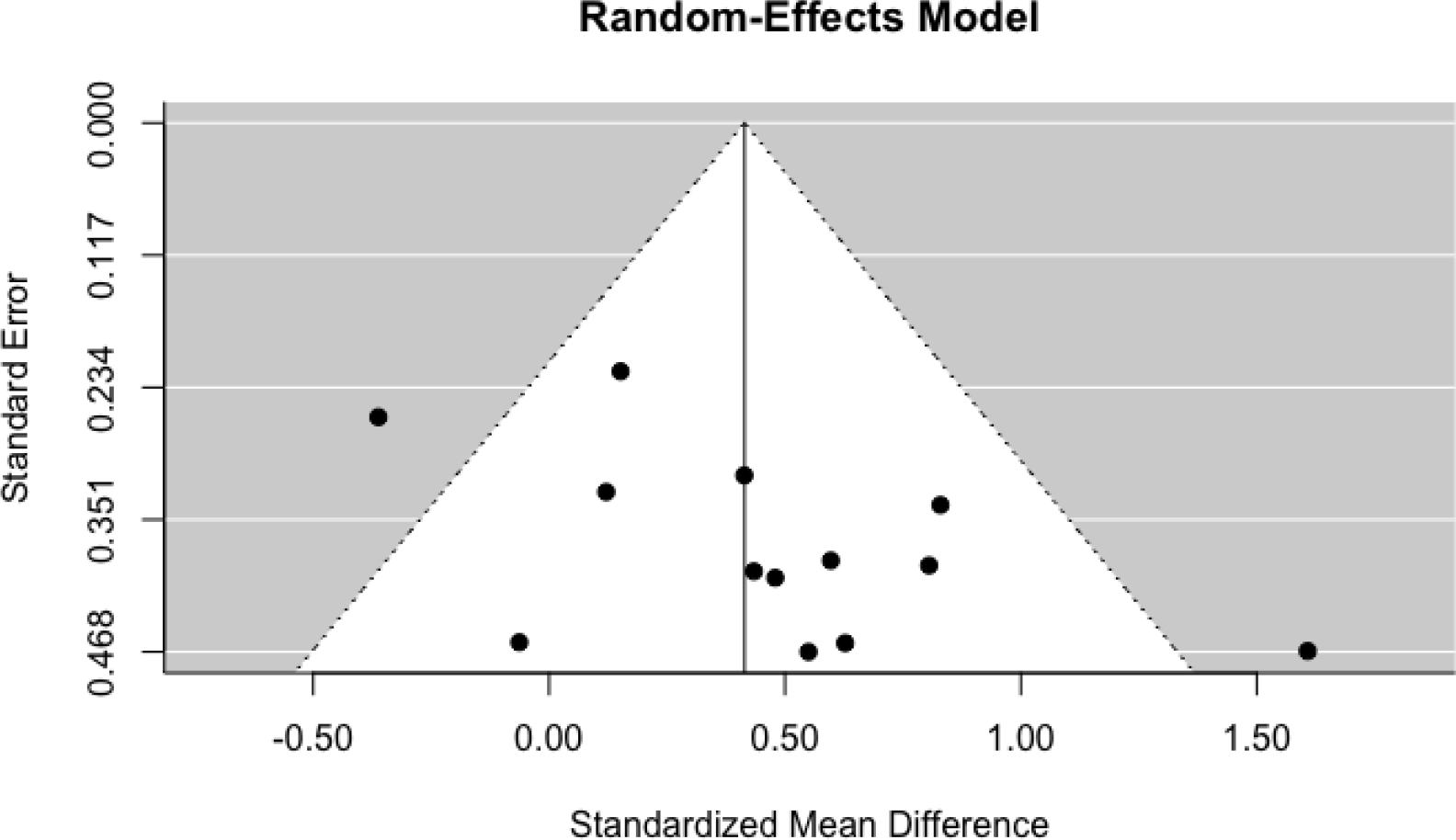
Funnel plot of publication bias for studies investigating pitch mean. The x-axis reports the effect size (Cohen's d) of the difference in pitch mean between ASD and comparison populations: positive values indicate higher mean pitch in ASD, while negative lower. The y-axis reports the standard error in each study. The white triangle represents an estimation of the real effect size distribution. The publication bias can be observed in the studies being organized on a diagonal line: higher standard error corresponding to bigger effect size.

*Pitch variability* indicates the magnitude of changes in pitch across the linguistic unit analysed (be it a phoneme, a word or a longer utterance). Pitch variability was investigated in 22 studies involving 398 participants with ASD and 337 comparison participants. 12 studies reported significant results, 11 indicating wider, one narrower and 10 no significant differences in pitch variability^3^. As all studies but two used pitch range, rarely adding measures of standard deviation and coefficient of variation, we based the meta-analysis on pitch range, introducing other measures only when range was not available.

The meta-analysis involved 17 studies, 320 participants with ASD and 275 comparison participants (see Figure 3). The overall estimated difference (Cohen's d) in pitch variability between the ASD and the comparison groups was 0.5 (95% CIs: 0.24 0.77, p=0.0002) with an overall variance (τ^2^) of 0.18 (95% CIs: 0.04 0.61). Much of the variance (I^2^: 60.18%, 95% CIs: 26.83 83.38) could not be reduced to random sample variability between studies (Q-stats=39.94, p=0.0008). However, neither task (estimate: 0.2, 95% CIs −0.15 0.55, p=0.27) nor language (estimate: −0.03, 95% CIs −0.12 0.05, p=0.42) could significantly explain the variance.

**Figure 3.**
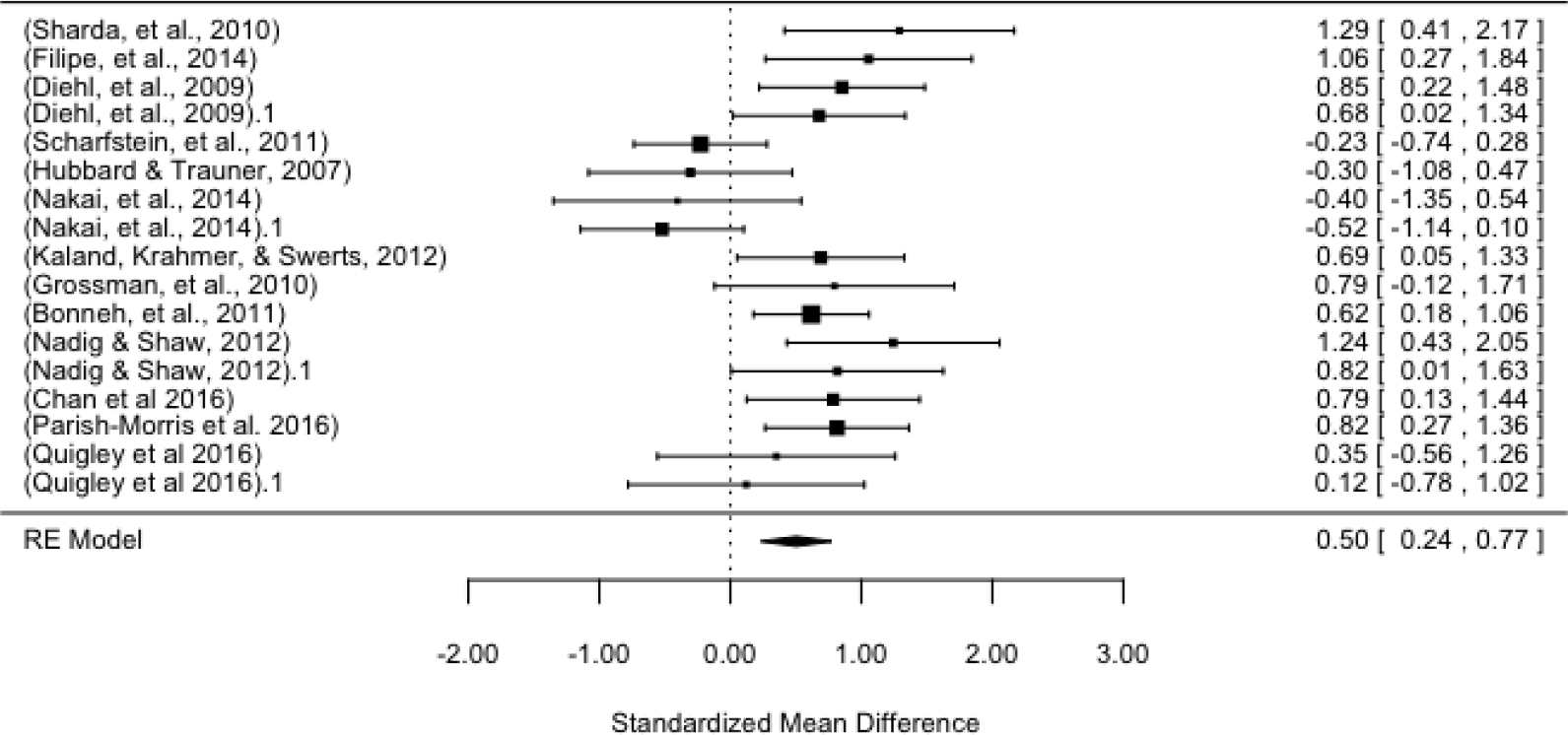
Forest plot of effect sizes (Cohen's d) in pitch range between the ASD and comparison populations. The x-axis reports the effect size (positive values indicate higher pitch variability in ASD, while negative lower) and the y-axis the studies for which statistical estimates of pitch mean were provided. The dotted vertical line indicates the null hypothesis (no difference between the populations).

There were no obvious outliers, nor any obvious publication bias (Kendall's τ=0.06, p=0.78; Figure 4). Indeed, of the 4 studies where statistical estimates were not available, 2 reported null findings and 2 included cases in which participants with ASD presented a wider pitch range, slightly reinforcing the hypothesis of a positive effect size.

**Figure 4.**
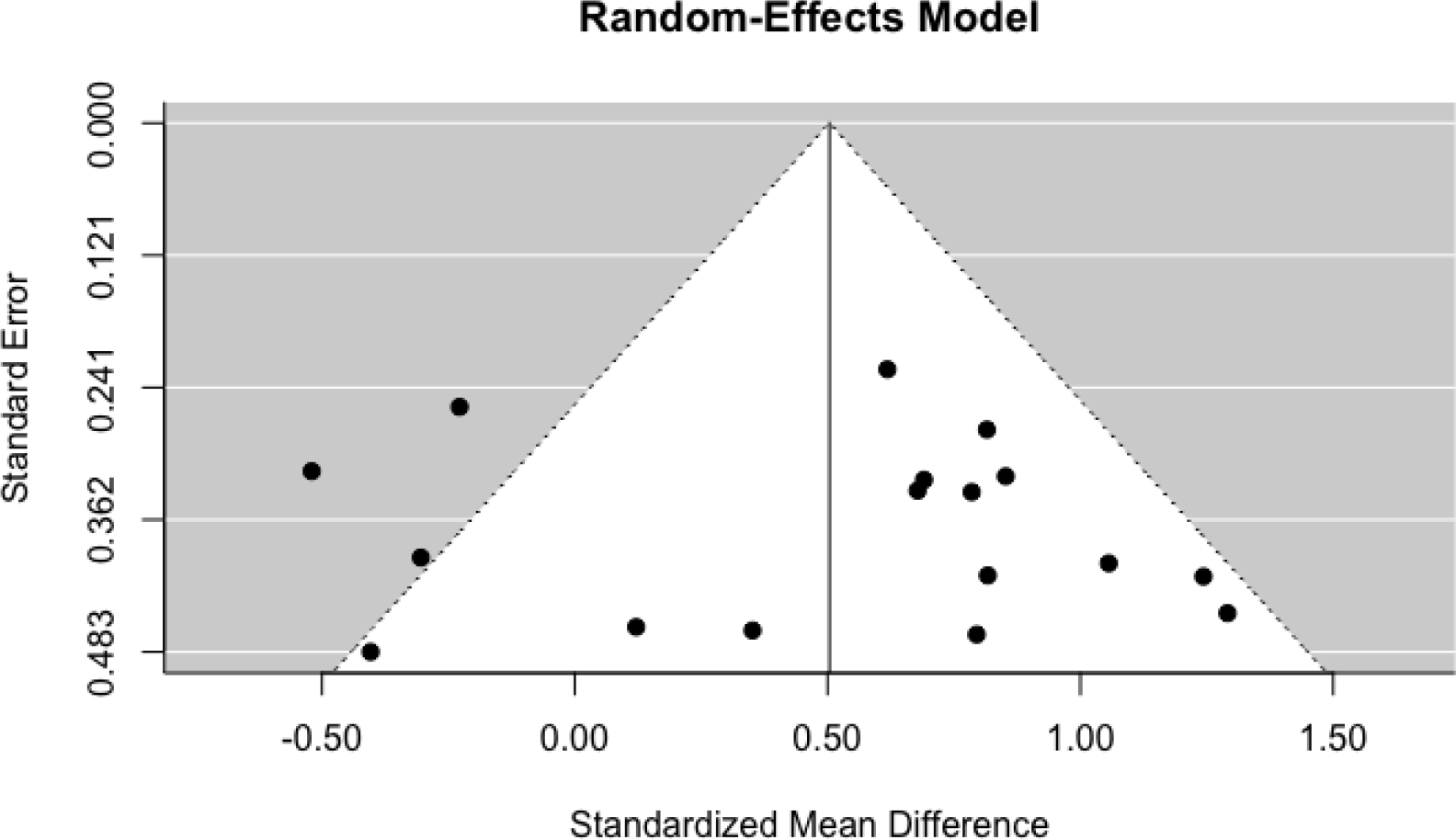
Funnel plot of publication bias for studies investigating pitch range. The x-axis reports the effect size (Cohen's d) of the difference in pitch mean between ASD and comparison populations: positive values indicate higher pitch variability in ASD, while negative lower. The y-axis reports the standard error in each study. The white triangle represents an estimation of the real effect size distribution.

*Pitch and severity of clinical features* were investigated in 5 studies (Table 2), which sought to relate quantitative measures of pitch measures to severity of clinical features as measured by the Autism Diagnostic Observation Schedule (ADOS, Lord, 2008) and the Autism Screening Questionnaire (ASQ, Dairoku, Senju, Hayashi, Tojo, & Ichikawa, 2004). Total ADOS scores were negatively related to the temporal trajectory of pitch. In particular, the steeper the slope of pitch change at the end of participants' speech turns, the lower the ADOS score (Bone, et al., 2014). However, null findings were reported in relation to pitch mean and range (Nadig & Shaw, 2012), and other temporal properties of pitch (Bone, et al., 2014). The communication sub-scale of the ADOS was found to correlate with pitch standard deviation in adolescents but not in children during narrative productions (Diehl, et al., 2009). Finally, pitch coefficient of variation was found to correlate negatively with ASQ Social Reciprocal Interaction, but not with total ASQ, Repetitive Behavior and Communication in children (Nakai, et al., 2014). As the direction of relation between pitch variability and clinical features seems to vary by study and no replication of any result is available, the current evidence is deemed inconclusive.

**Table 2.**
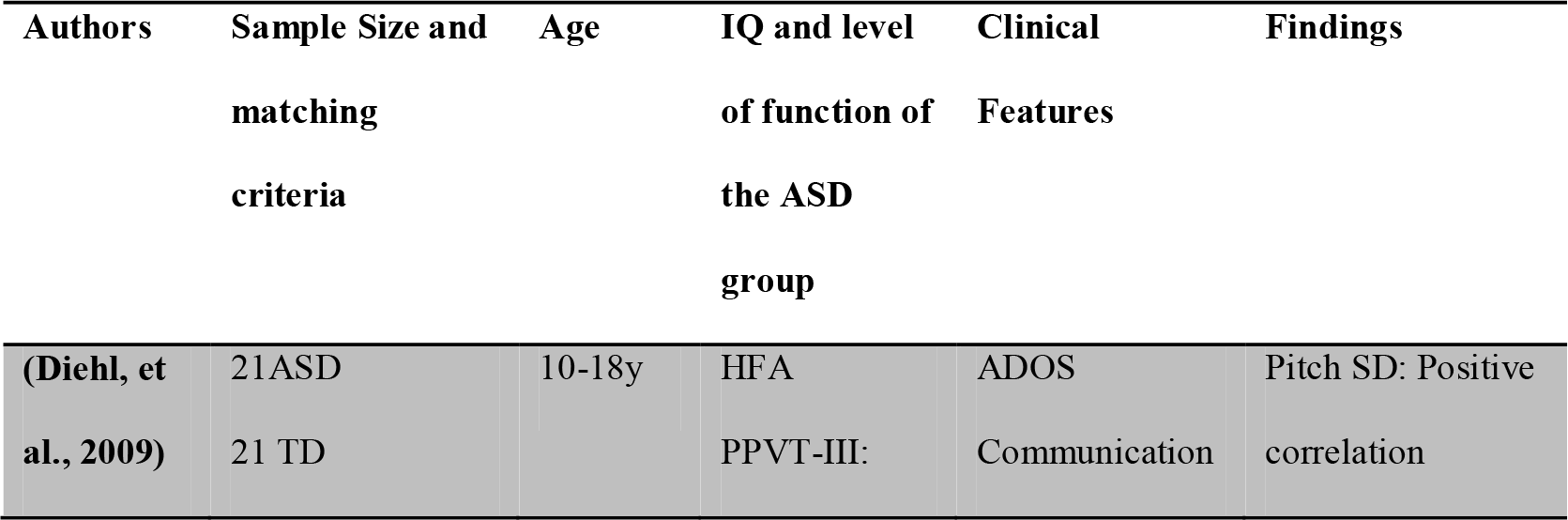
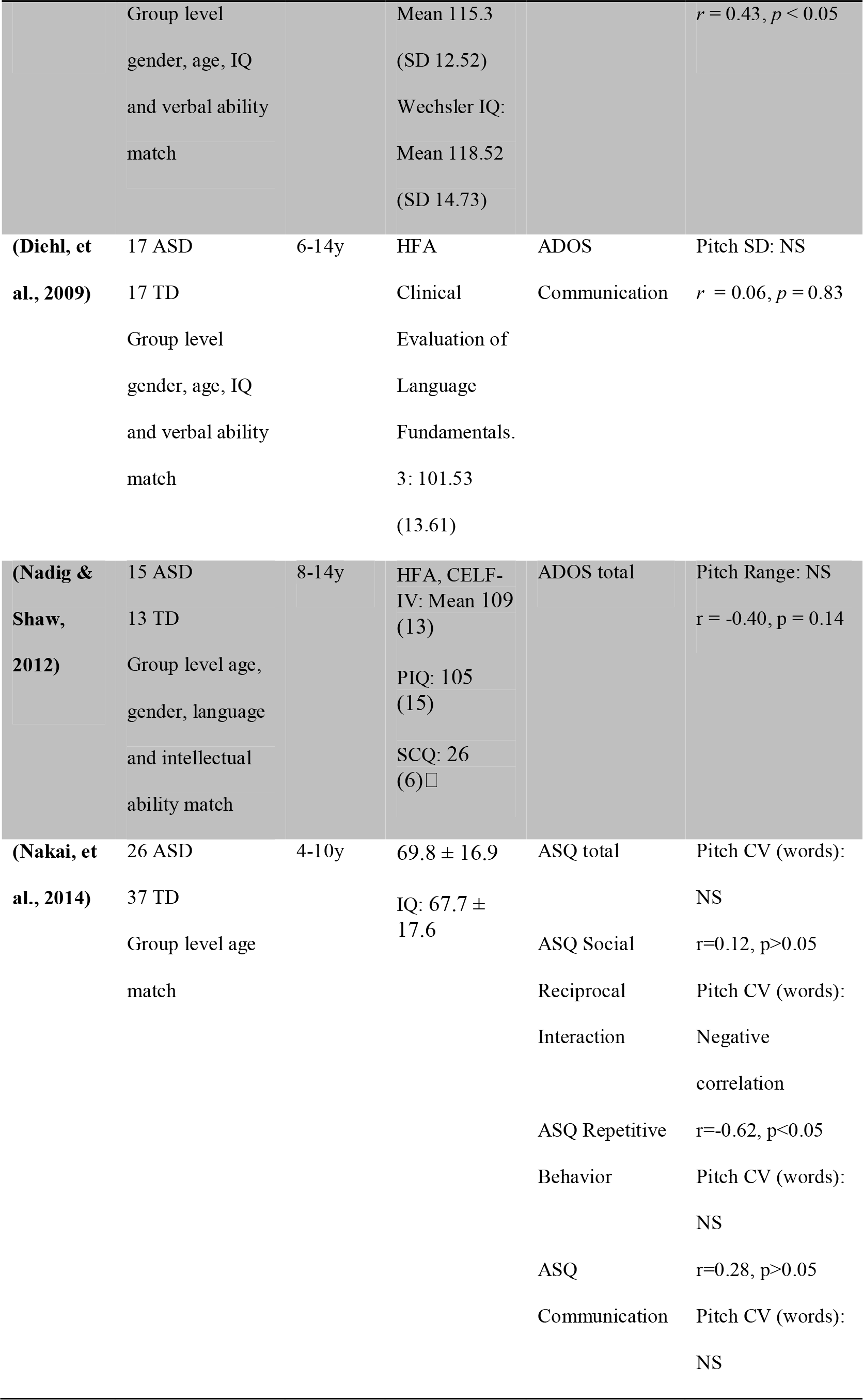
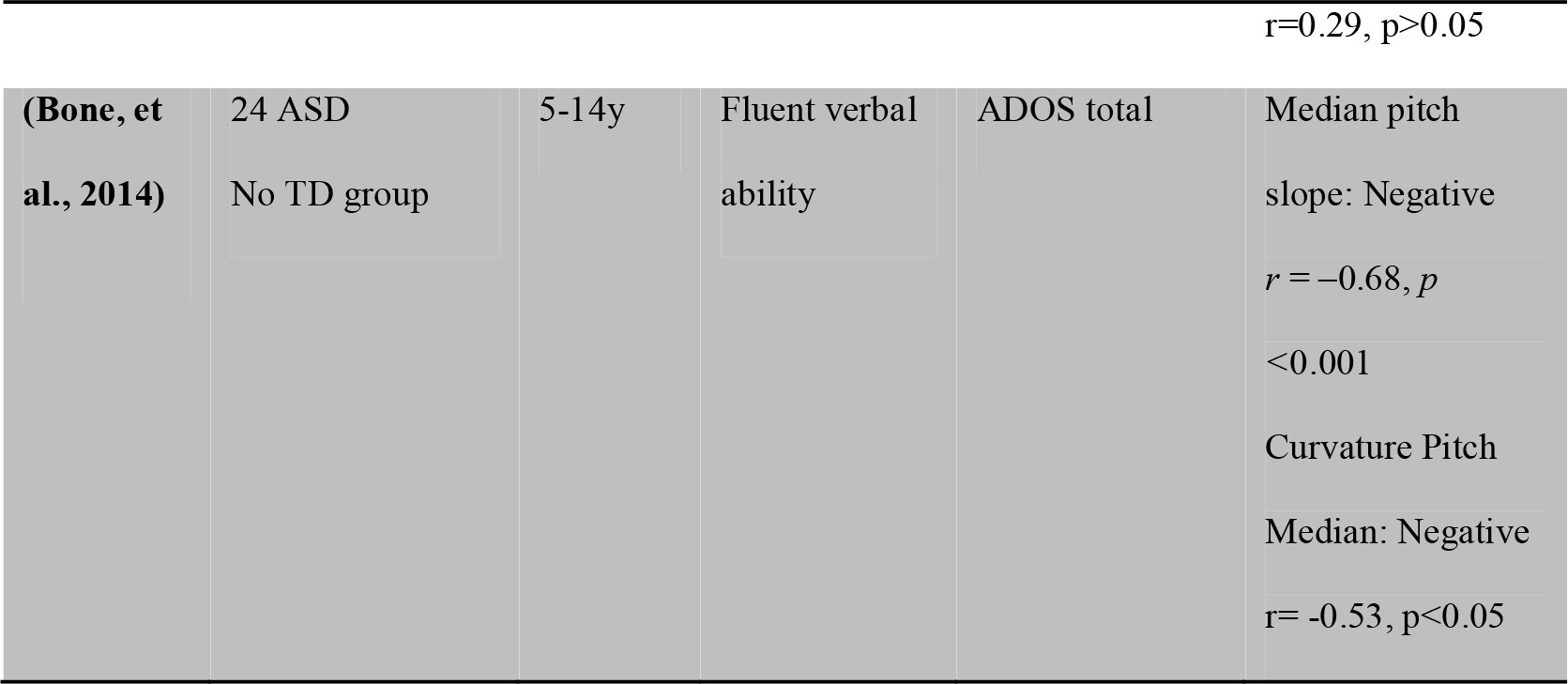
*Relations between acoustic measures and severity of clinical features*.

While anecdotal and qualitative reports clearly indicate a difference in the use of pitch in ASD, the acoustic evidence is more uncertain, with little replication, and a high number of non-significant or contradictory findings. Even taking at face value the two meta-analytic effect sizes, it should be noted that an estimated difference of Cohen's d 0.4 to 0.5 is a small difference. Indeed, if we were to use these statistical estimates to guess whether any given voice belongs to a participant with ASD or to a comparison one, we would only be right about 61–64% of the time, an insufficient level of accuracy to justify its use as a potential marker (Ellis, 2010).

### 3.2 Intensity

Intensity or loudness is a measure of the energy carried by a sound wave and is important for making speech intelligible and for expressing emotions. 8 studies have investigated intensity through quantitative measures (Table 3).

**Table 3.**
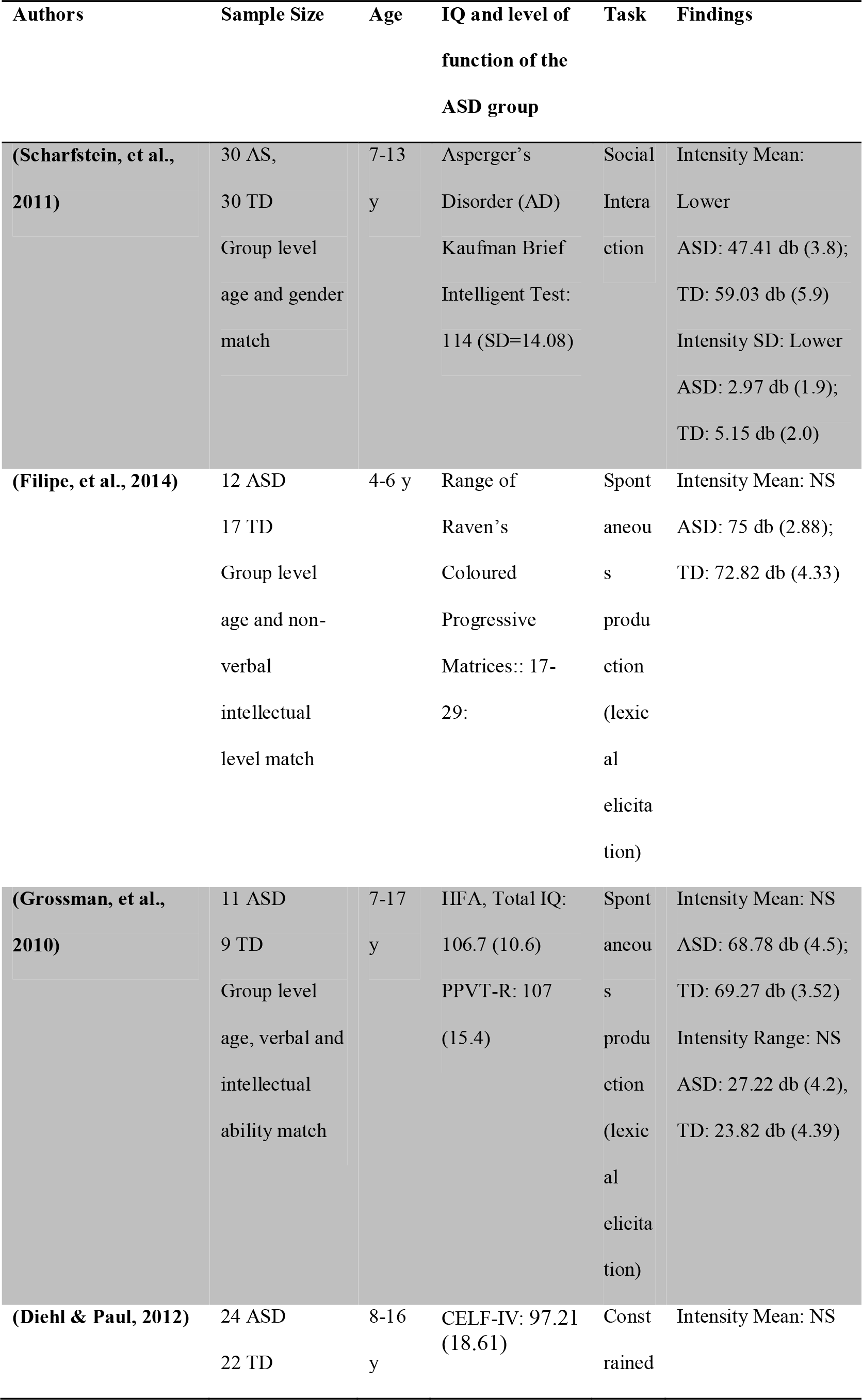
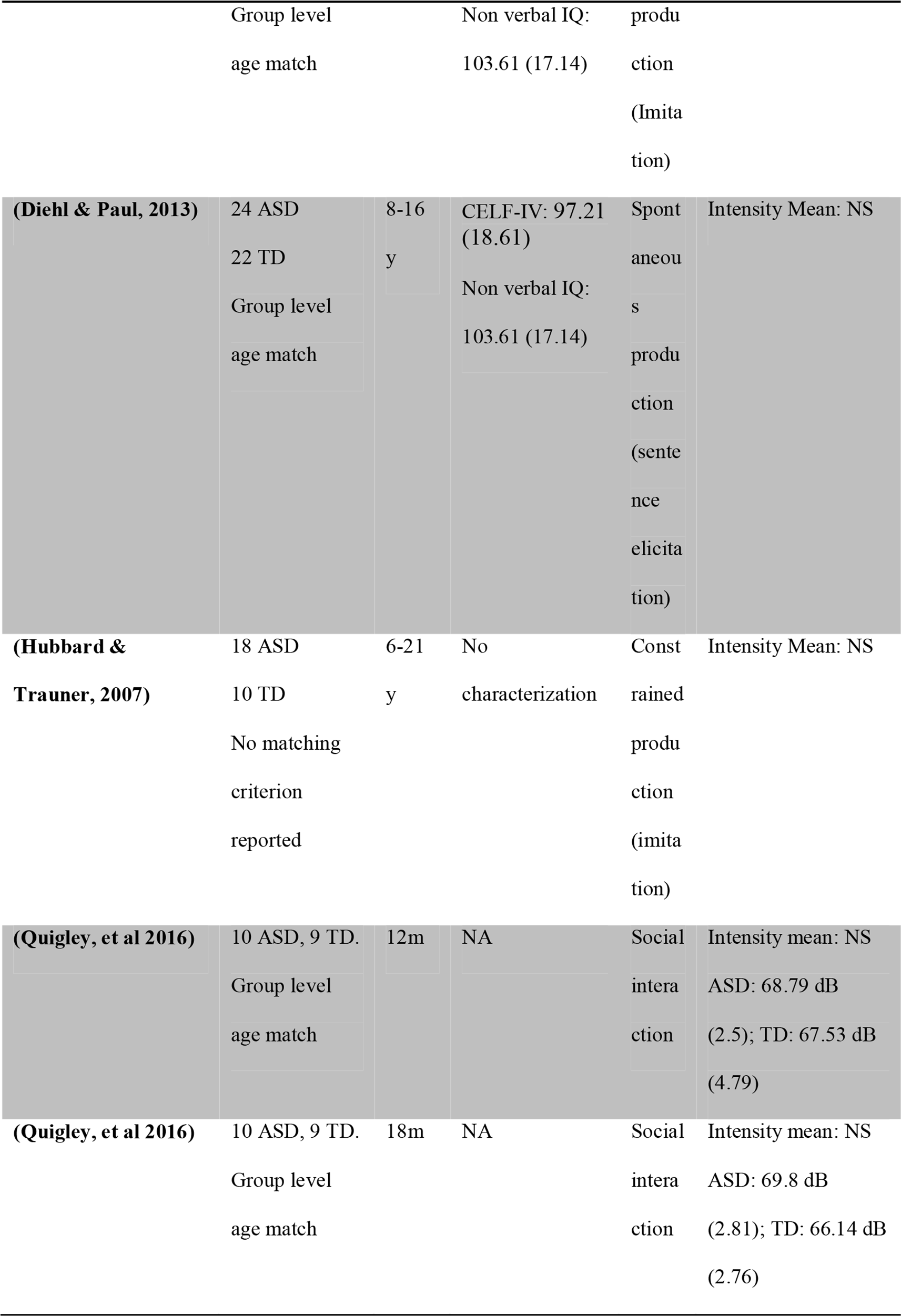
*Studies involving acoustic measures of intensity in ASD*.

| Authors | Sample Size | Age | IQ and level of function of the ASD group | Task | Findings |
| --- | --- | --- | --- | --- | --- |
| (Scharfstein, et al., 2011) | 30 AS,<br>30 TD<br>Group level<br>age and gender<br>match | 7-13<br>y | Asperger's<br>Disorder (AD)<br>Kaufman Brief<br>Intelligent Test:<br>114 (SD=14.08) | Social<br>Intera<br>ction | Intensity Mean:<br>Lower<br>ASD: 47.41 db (3.8);<br>TD: 59.03 db (5.9)<br>Intensity SD: Lower<br>ASD: 2.97 db (1.9);<br>TD: 5.15 db (2.0) |
| (Filipe, et al., 2014) | 12 ASD<br>17 TD<br>Group level<br>age and non-verbal<br>intellectual<br>level match | 4-6 y | Range of<br>Raven's<br>Coloured<br>Progressive<br>Matrices:: 17-29: | Spont<br>aneou<br>s<br>produ<br>ction<br>(lexic<br>al<br>elicit<br>ation) | Intensity Mean: NS<br>ASD: 75 db (2.88);<br>TD: 72.82 db (4.33) |
| (Grossman, et al., 2010) | 11 ASD<br>9 TD<br>Group level<br>age, verbal and<br>intellectual<br>ability match | 7-17<br>y | HFA, Total IQ:<br>106.7 (10.6)<br>PPVT-R: 107<br>(15.4) | Spont<br>aneou<br>s<br>produ<br>ction<br>(lexic<br>al<br>elicit<br>ation) | Intensity Mean: NS<br>ASD: 68.78 db (4.5);<br>TD: 69.27 db (3.52)<br>Intensity Range: NS<br>ASD: 27.22 db (4.2),<br>TD: 23.82 db (4.39) |
| (Diehl & Paul, 2012) | 24 ASD<br>22 TD | 8-16<br>y | CELF-IV: 97.21<br>(18.61) | Const<br>rained | Intensity Mean: NS |
|  | Group level |  | Non verbal IQ: | produ |  |
|  | age match |  | 103.61 (17.14) | ction |  |
|  |  |  |  | (Imita |  |
|  |  |  |  | tion) |  |
| **(Diehl & Paul, 2013)** | 24 ASD | 8-16 | CELF-IV: 97.21 (18.61) | Spont | Intensity Mean: NS |
|  | 22 TD | y |  | aneou |  |
|  | Group level |  | Non verbal IQ: | s |  |
|  | age match |  | 103.61 (17.14) | produ |  |
|  |  |  |  | ction |  |
|  |  |  |  | (sente |  |
|  |  |  |  | nce |  |
|  |  |  |  | elicit |  |
|  |  |  |  | ation) |  |
| **(Hubbard & Trauner, 2007)** | 18 ASD | 6-21 | No | Const | Intensity Mean: NS |
|  | 10 TD | y | characterization | rained |  |
|  | No matching |  |  | produ |  |
|  | criterion |  |  | ction |  |
|  | reported |  |  | (imita |  |
|  |  |  |  | tion) |  |
| **(Quigley, et al 2016)** | 10 ASD, 9 TD. | 12m | NA | Social | Intensity mean: NS |
|  | Group level |  |  | intera |  |
|  | age match |  |  | ction |  |
|  |  |  |  |  | ASD: 68.79 dB |
|  |  |  |  |  | (2.5); TD: 67.53 dB |
|  |  |  |  |  | (4.79) |
| **(Quigley, et al 2016)** | 10 ASD, 9 TD. | 18m | NA | Social | Intensity mean: NS |
|  | Group level |  |  | intera |  |
|  | age match |  |  | ction |  |
|  |  |  |  |  | ASD: 69.8 dB |
|  |  |  |  |  | (2.81); TD: 66.14 dB |
|  |  |  |  |  | (2.76) |

*Intensity Mean* was available for 8 studies (105 ASD and 97 comparison participants), one with significantly lower intensity for ASD and the others with null findings (Filipe, et al., 2014; Grossman, et al., 2010; Scharfstein, et al., 2011).

*Intensity variability* was available for 2 studies involving 41 ASD and 39 comparison participants. One study reported lower variability, and the other null findings.

Finally, one study attempted to relate intensity measures and severity of clinical features (ADOS total score): No significant correlation was found for ADOS and the temporal profiles of intensity, such as slope and curvature (Bone, et al., 2014).

In summary, there is not enough acoustic evidence to support the impression of atypical voice intensity in ASD. It should be noted that acoustic measures of intensity are highly dependent on the relative positions of microphone and speakers, as well as to changes in angle and distance through the vocal production and therefore highly prone to external artifacts. Intensity measures should therefore be assessed with caution.

### 3.3. Duration, speech rate and pauses

Duration is measured as length in seconds, and has been applied to full utterances, lexical items (words) and syllables (often distinguishing between stressed and unstressed syllables). A related duration measure, speech rate, is measured as estimated syllables per second, number of pauses, length of pauses and voiced duration. 19 studies employed acoustic descriptors of duration, pauses and speech rate (see Table 4).

**Table 4.**
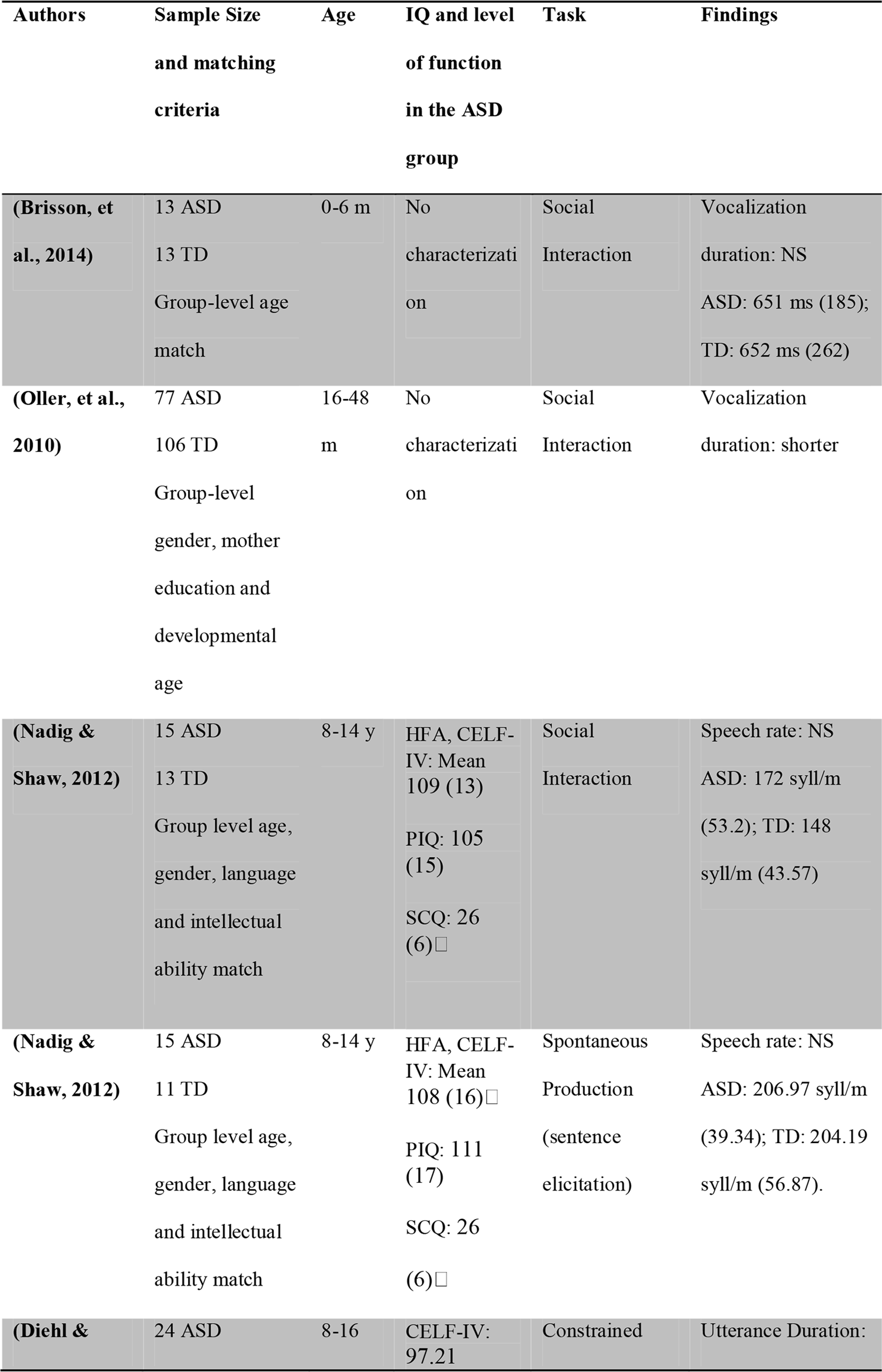
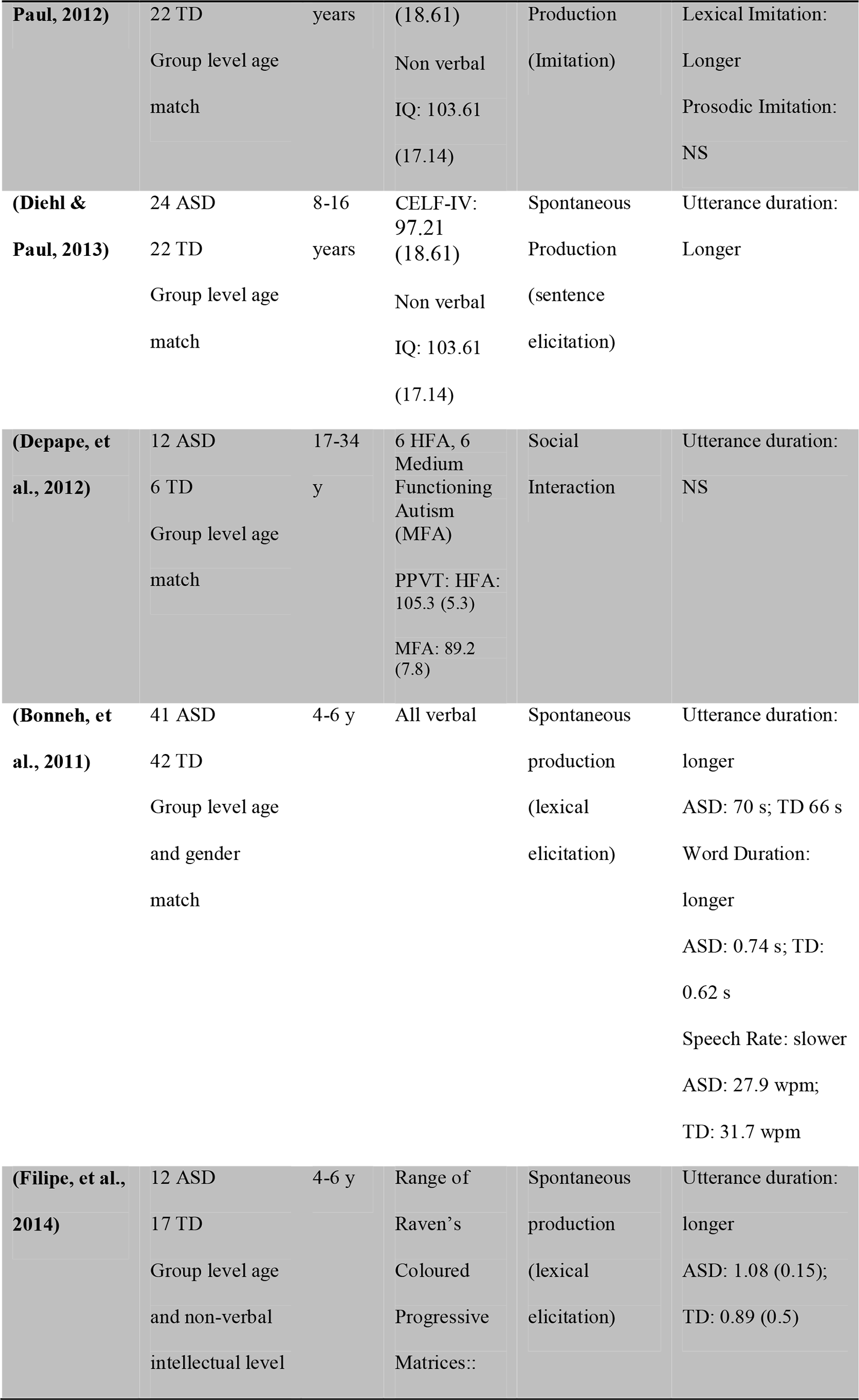
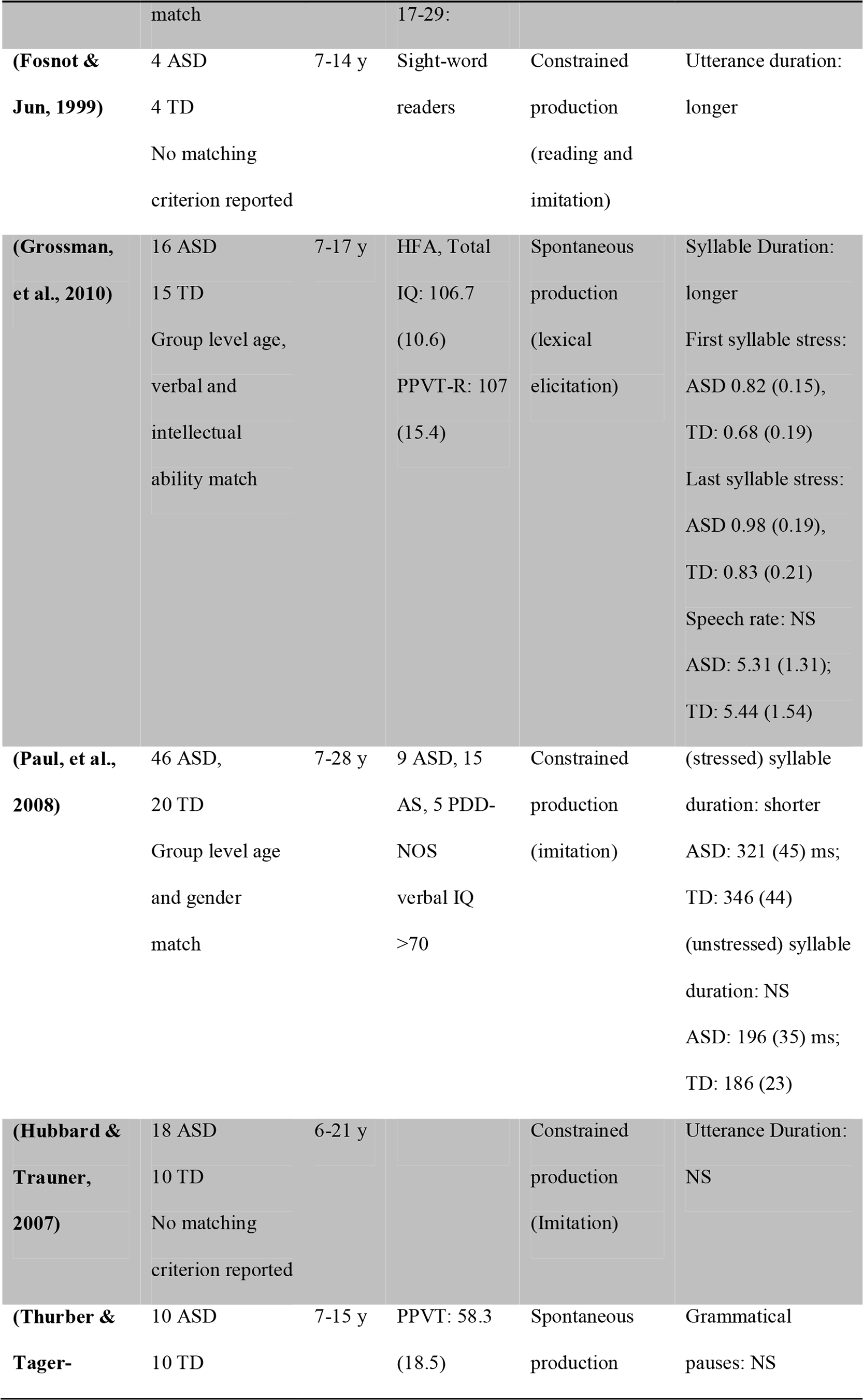
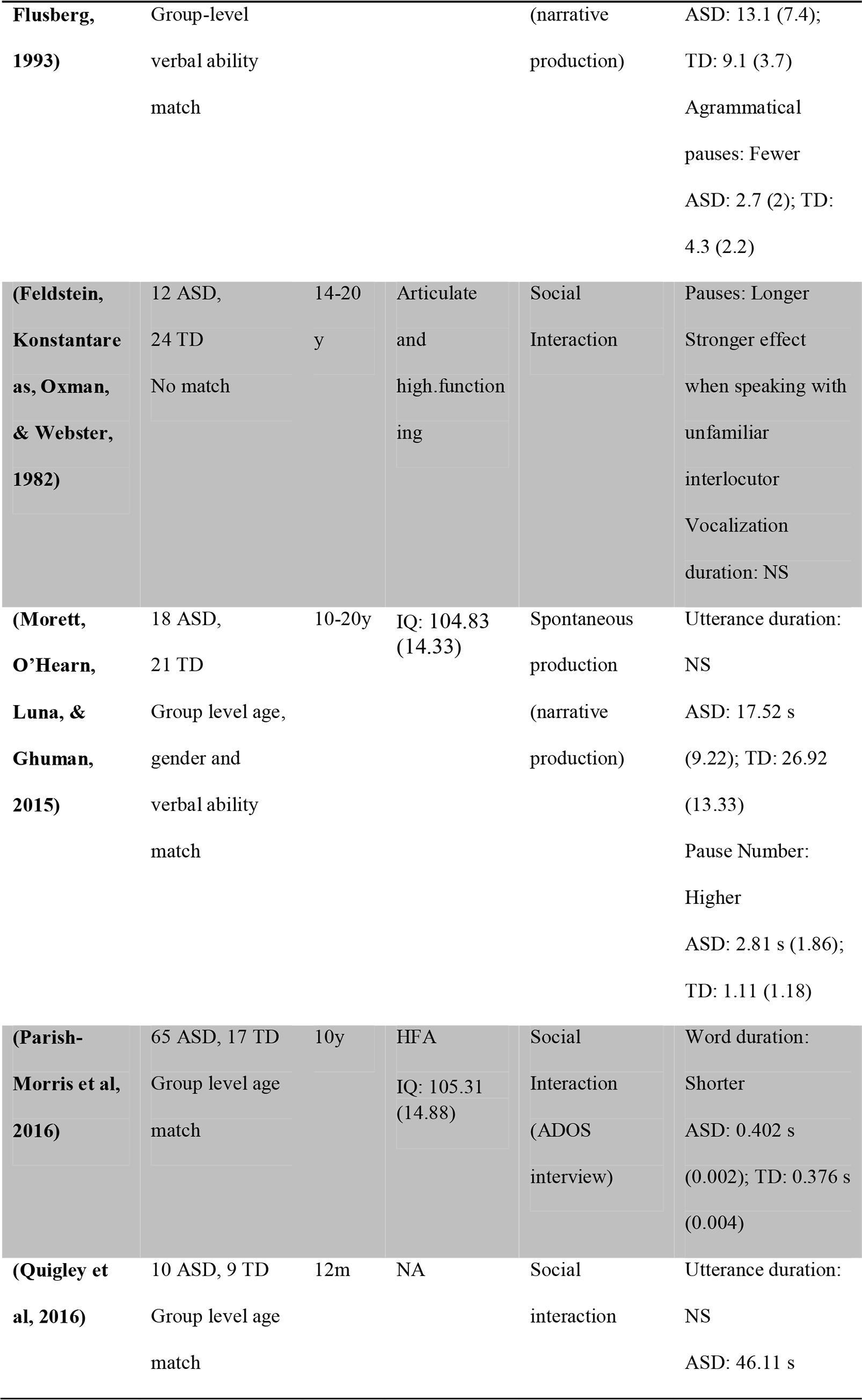
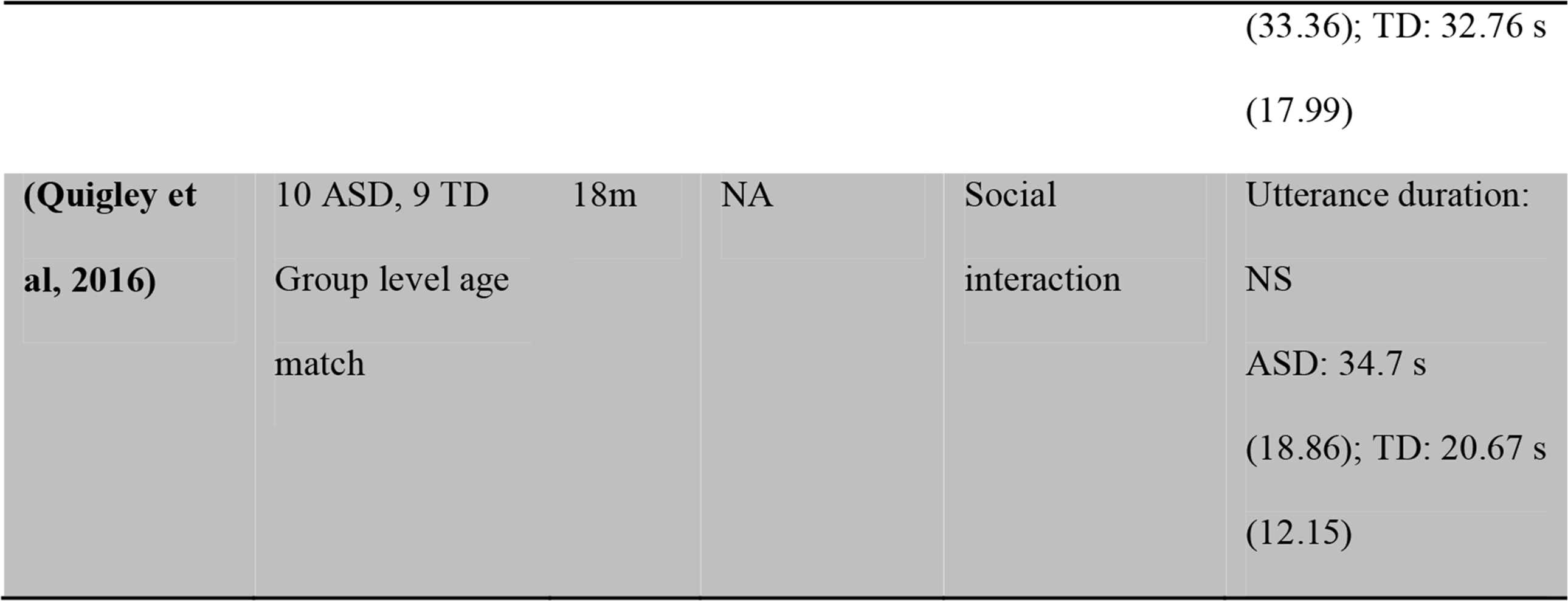
*Studies involving acoustic measures of intensity in ASD*.

Out of 15 studies involving duration measures 7 reported longer duration, 6 reported no differences between groups and 1 shorter duration in ASD. Out of 4 studies investigating speech rate, 3 reported null findings and 1 found slower speech rate in ASD. Out of 2 studies focusing on syllable duration, one reports longer duration for stressed syllables in ASD, whereas the other reports shorter duration for stressed syllables and no differences for unstressed syllables. Out of 3 studies measuring speech pauses, 1 finds longer pauses, 1 no difference in grammatically motivated pauses, but fewer pragmatically motivated ones and the third a higher number of pauses. Two studies investigated the relation between speech rate and severity of clinical features in terms of ADOS total scores), but found no significant correlations (Bone, et al., 2014; Nadig & Shaw, 2012). In sum, not enough statistical estimates were reported to allow for meta-analyses and the findings do not seem conclusive.

### 3.4. Voice Quality

Voice quality covers a large variety of features, which do not overlap between studies. Hoarseness, breathiness and creaky voice are often attributed to imperfect control of the vocal fold vibrations that produce speech and have been quantified as irregularities in pitch (jitter) and intensity (shimmer), or as low harmonic to noise ratio (relation between periodic and aperiodic sound waves) (Tsanas, Little, McSharry, & Ramig, 2011). More generic definitions of dysphonia, or voice perturbation, rely on cepstral analyses, which involve a further frequency decomposition of the pitch signal, that is, the frequency of changes in frequency (Maryn, Roy, De Bodt, Van Cauwenberge, & Corthals, 2009). Analyses of voice quality are particularly challenging and difficult to compare across studies because of a lack of established standards: they rely on the choice of several parameters, and the results change greatly if applied to prolonged phonations (held vowels), or continuous speech (Laver, Hiller, & Beck, 1992; Orlikoff & Kahane, 1991).

So far only one published study has investigated acoustic measures of voice quality in ASD: children with ASD were shown to have more jitter and jitter variability, as well as less harmonic to noise ratio, and no differences in shimmer or cepstral peak prominence (Bone, et al., 2014). However, a series of unpublished conference papers point to breathiness (Boucher, Andrianopoulos, & Velleman, 2010; Wallace et al., 2008), tremors (Wallace, et al., 2008), and task-and vowel-dependent low jitter and low shimmer (Boucher, Andrianopoulos, Velleman, & Pecora, 2009).

One study investigated the relation between ADOS total scores and voice quality, highlighting positive correlations with jitter and harmonics to noise ratio variability, and negative ones with levels of Harmonic to Noise Ratio (Bone, et al., 2014). Notice that since the only published study mentioned here is already fully reported in previous tables, we have not produced a dedicated table for studies on voice quality.

In summary, while a distinctive voice quality has been reported in ASD since the very early days of the diagnosis, quantitative evidence is extremely sparse. While potentially promising, the existing studies use non-overlapping measures, making it difficult to assess the generality of the patterns observed.

## 4. Results: From Acoustic Patterns to Diagnosis (multivariate machine learning studies)

The previous section reviewed studies identifying differences in acoustic patterns produced by ASD and comparison samples, one feature at a time. In this section we review a second set of 15 studies (see Table 5), which present an alternative approach: multivariate machine-learning (Bishop, 2006; Hastie, Tibshirani, & Friedman, 2009). Briefly, multivariate machine learning differs from traditional univariate approaches in three respects. First, the research question is reversed. Univariate approaches ask whether there is a statistically significant difference between two distinct populations (independent variable) with respect to some measure (dependent variable). Machine learning approaches seek to determine whether the data contains enough information to accurately separate the two populations. Second, a multivariate approach enters multiple data features simultaneously into the analysis, including a wider variety of features than normally treated in their simple univariate form (such as more detailed spectral and cepstral features, see par. 3.4). Third, the goal is not to identify the statistical model that best separates the populations from which the data has been obtained, but to identify the model that best generalizes to new data (e.g., generalize from a training to a test set of data, see Yarkoni & Westfall, 2016).

Multivariate machine learning studies typically involve processes of 1) feature extraction, 2) feature selection and 3) classification (e.g., presence of diagnosis) or score prediction (e.g., severity of clinical features), the latter two often undergoing a process of 4) validation.

The first process involves extraction of acoustic features from vocal recordings. Most studies use summary statistics discussed in the earlier section (mean and standard deviation of acoustic features), but they often include additional measures, such as non-linear descriptive statistics. Traditional summary statistics cannot adequately capture the non-stationary nature of the speech signal; for example, the mean and the standard deviation of pitch often change over a speech event (Jiang, Zhang, & McGilligan, 2006). In contrast, time-aware measures - such as slope analysis, recurrence quantification analysis, Teager-Kaiser energy operator and fractal analyses - quantify the degree to which acoustic patterns change or are repeated in time (cf. Table 5. For detailed and technical descriptions of these methods, cf. Bone, et al., 2014; Kiss, van Santen, Prud'hommeaux, & Black, 2012; Marwan, Carmen Romano, Thiel, & Kurths, 2007; Riley, Bonnette, Kuznetsov, Wallot, & Gao, 2012; Tsanas, et al., 2011; Weed & Fusaroli, submitted). Finally, most studies expand the range of measures, by further quantifying formants, spectral and cepstral properties of the speech signal (cf. Table 5, for a more detailed treatment of these measures cf. the referred papers and Eadie & Doyle, 2005). Feature extraction is a largely automated process, but it often relies on basic manual pre-processing of the data: evaluation of background noise, isolation of the utterances, sometimes time-coding of the single words (e.g. Nakai et al 2014). However, it is still unclear how much hand-coding is theoretically necessary and promising automated techniques are being developed to replace it (e.g. Miro et al 2012; Xanguera et al. 2014).

As the first process very often generates a large number of acoustic features, the second process deals with identifying amongst them a minimal set of maximally informative features. A popular rule of thumb suggests that the feature selection process should select a number of features inferior to a tenth of the number of independent data points in the dataset, but different algorithms can deal with different ratios of features to data points. The third process involves the use of the selected features to construct a statistical model maximally distinguishing the target groups of interest (for detailed introductions to these topics, cf. Bishop, 2006; Hastie, et al.,2009) or most accurately predicting a score (e.g. severity of a given clinical feature).

Since the goal of machine learning procedures is not simply to explain the current data but to create models that generalize to new data, feature selection and classification are often validated (or cross-validated, (for details, cf. Rodriguez, Perez, & Lozano, 2010), for details). Validation involves the division of the dataset into training and test sets. The statistical models are fit to the training set and their explanatory power assessed on the test set.

The characteristics and findings of the multi-variate machine-learning studies are reported in Table 5. For a more detailed overview of how the different studies reviewed implement feature selection, classification and validation, see Supplementary Material S1.^4^

**Table 5.**
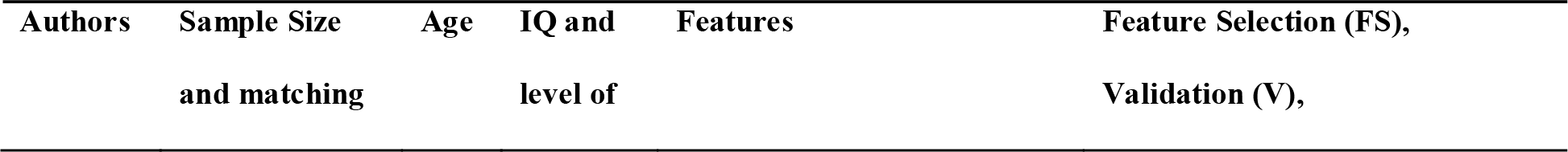
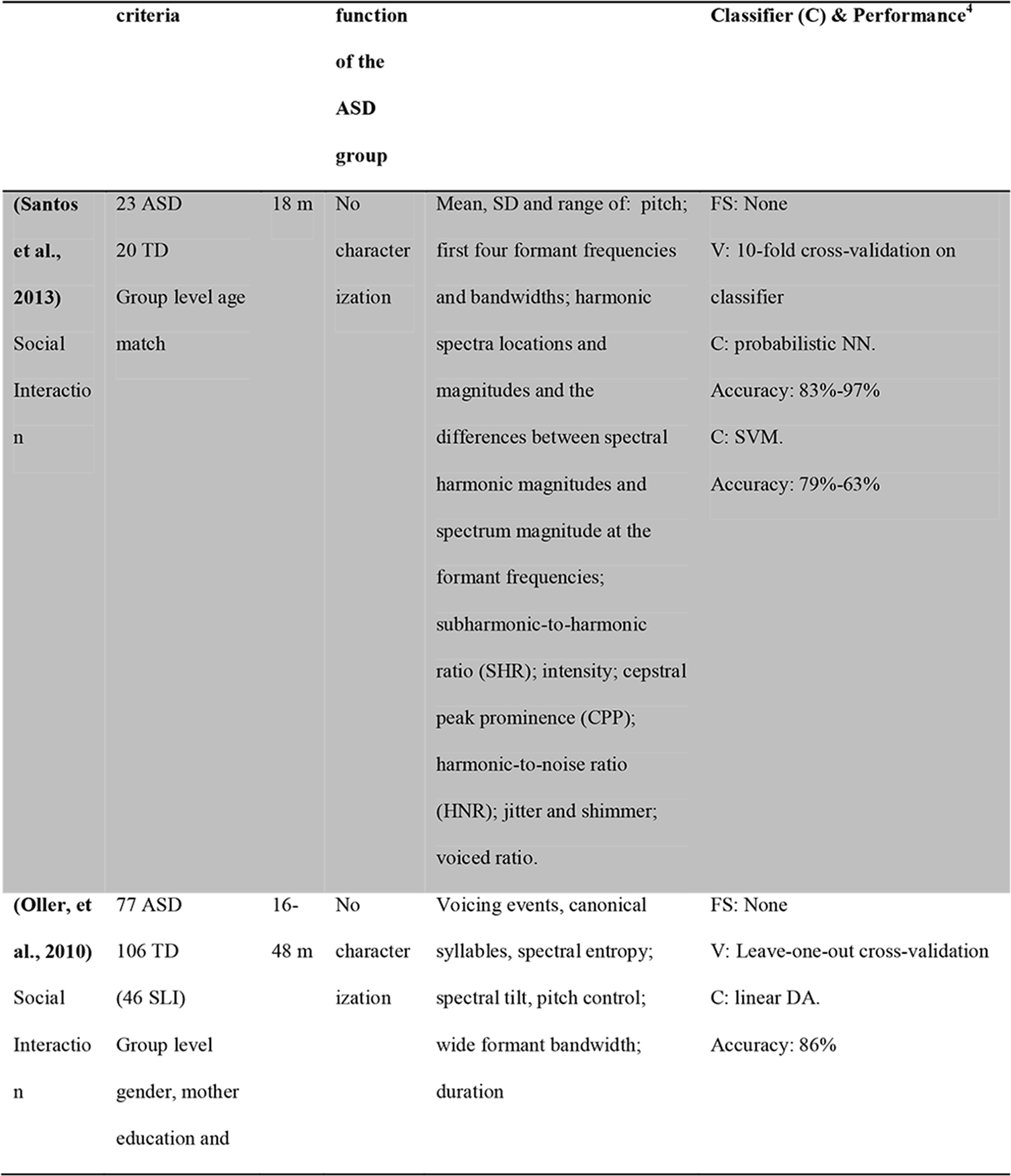
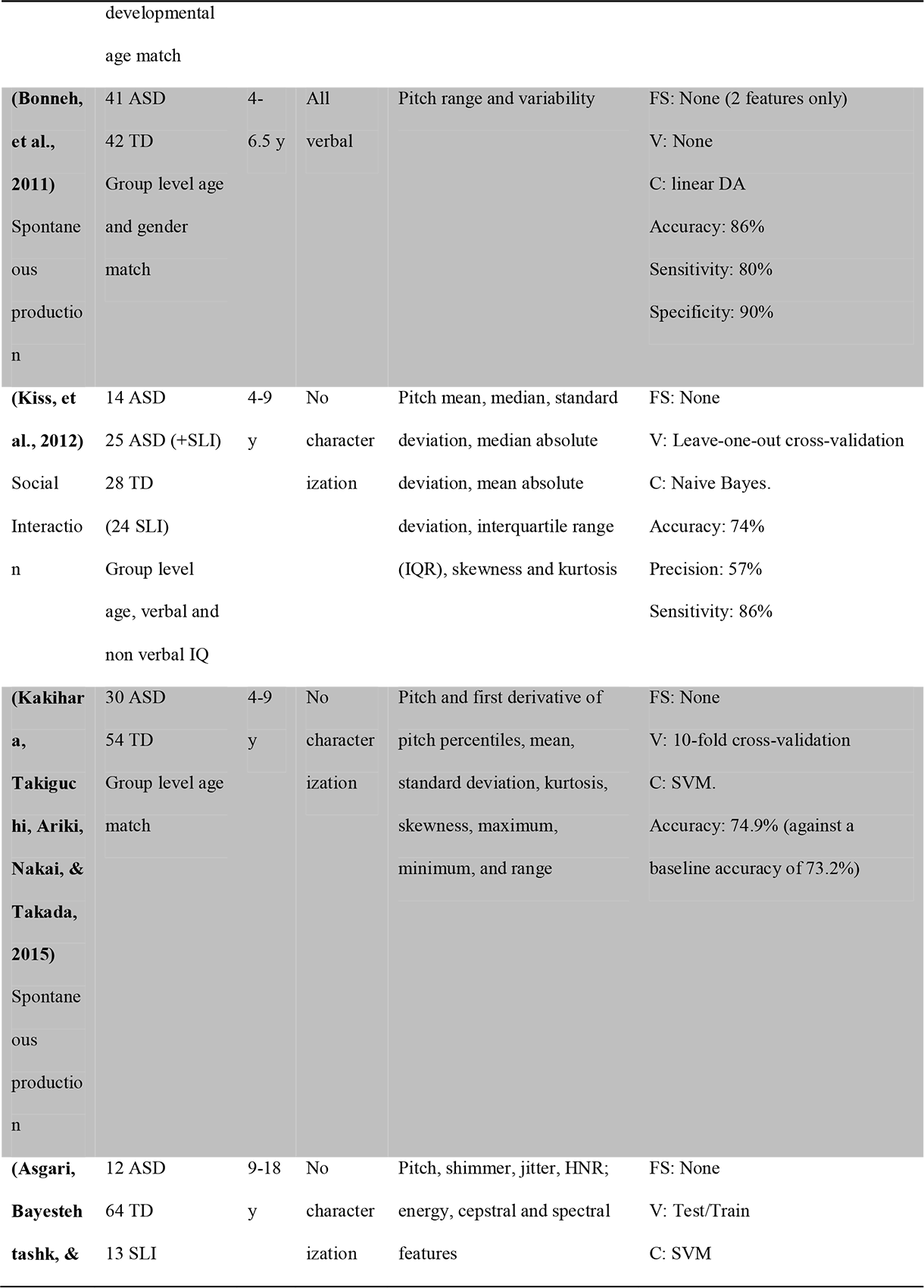
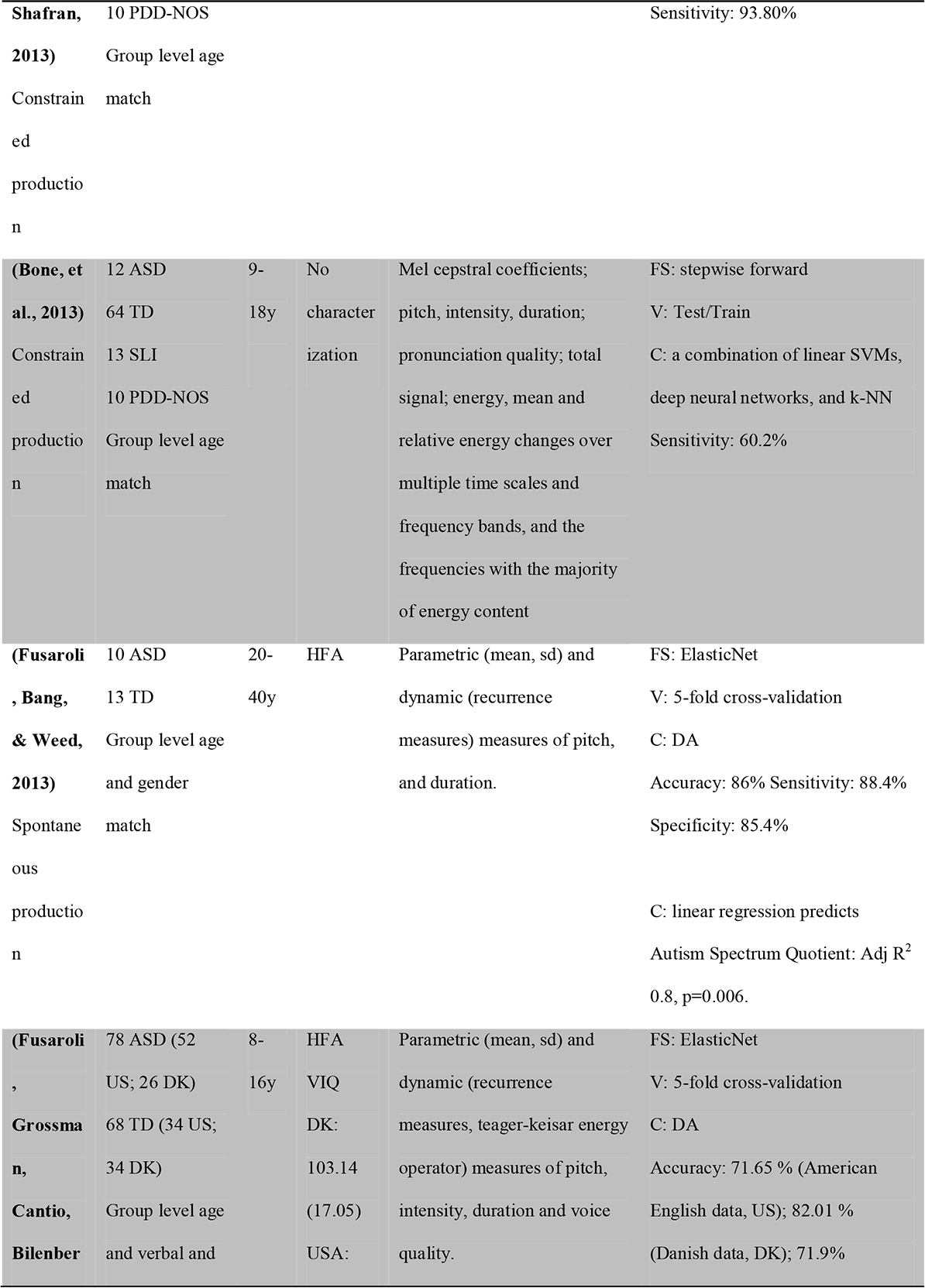
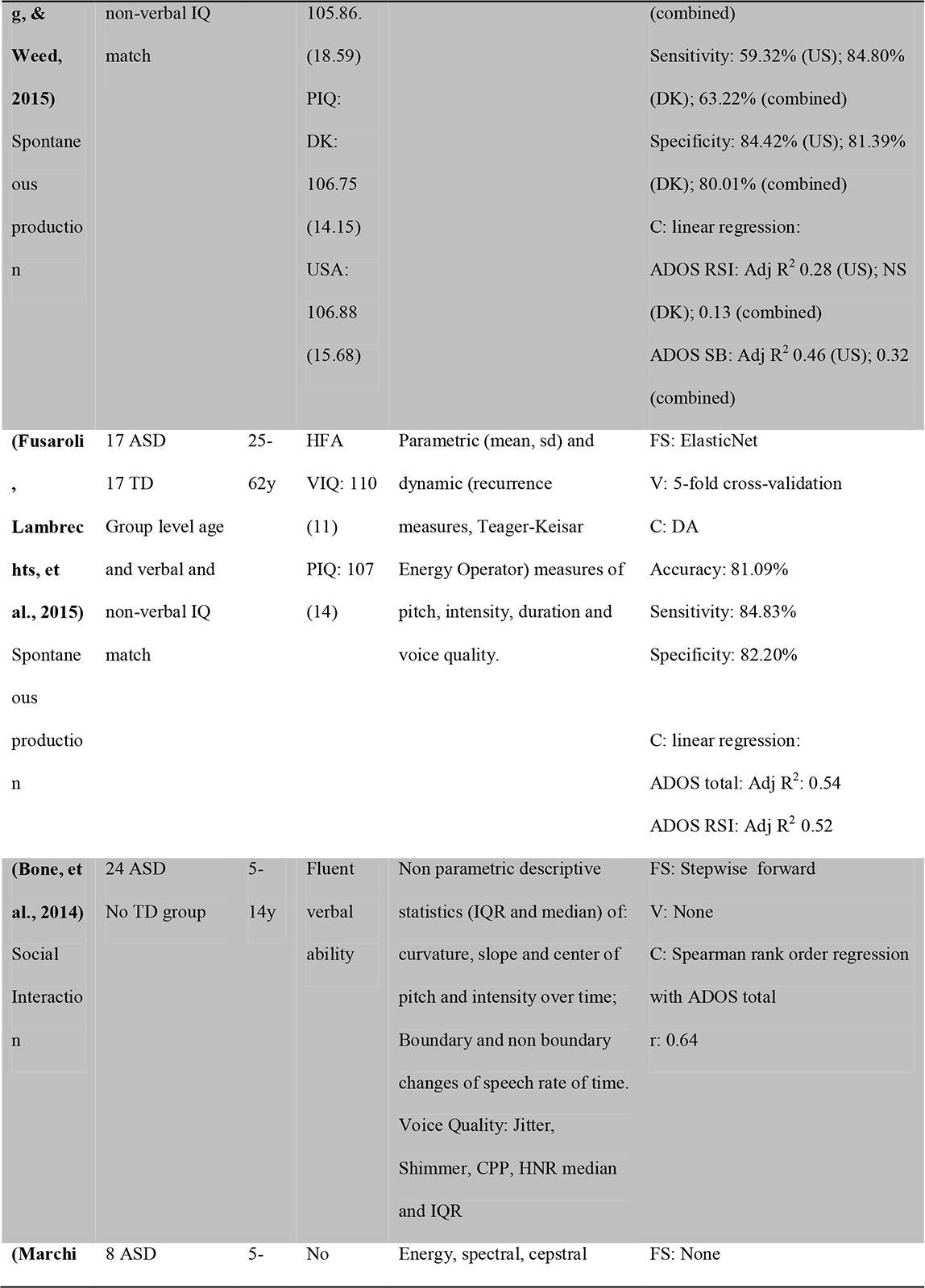
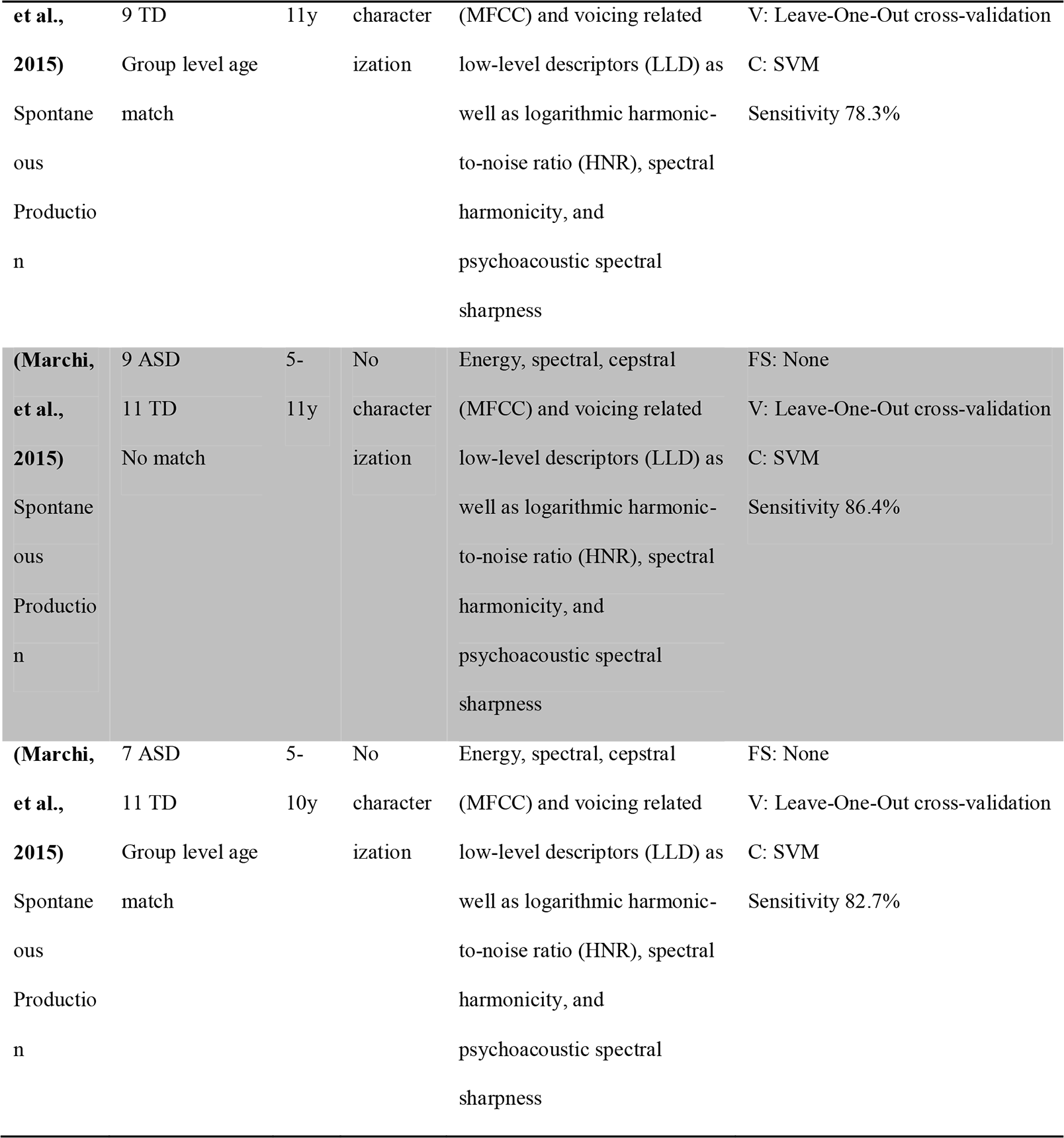
Reconstructing Diagnosis from Voice Patterns. An overviewdevelopmental age match

While simple measures of pitch were the most commonly employed, no single feature was used in all, or even in the majority of the studies. Analogously no single feature selection, classification algorithm or validation process was employed in a majority of studies. In terms of results, all but one multivariate machine-learning study reported accuracies well above 70% and up to 96%^5^. A more precise overview of the sensitivities and specificities of the algorithms, when it was possible to reconstruct them and their uncertainty, is presented in Figures 5 and 6. The average sensitivity was 80% (with one study indistinguishable from chance) and the average specificity was 85.1% (with all studies above chance).

**Figure 5.**
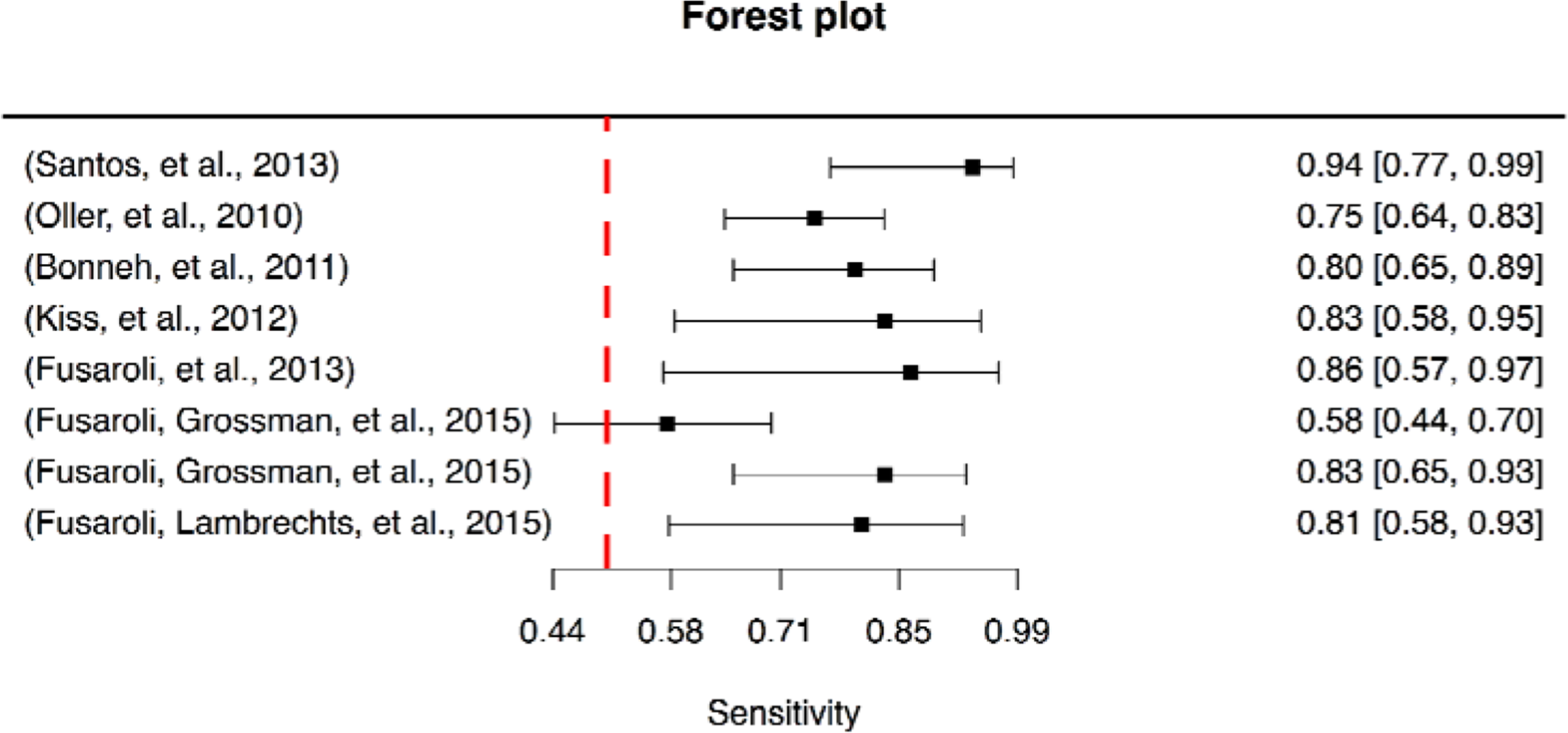
Forest plot of the algorithms' sensitivities in automatically discriminating between the ASD and comparison populations. The x-axis reports the sensitivity and the y-axis the studies for which it was possible to reconstruct the confidence intervals of sensitivity. The dotted line indicates sensitivity at chance level, that is, 50%.

**Figure 6.**
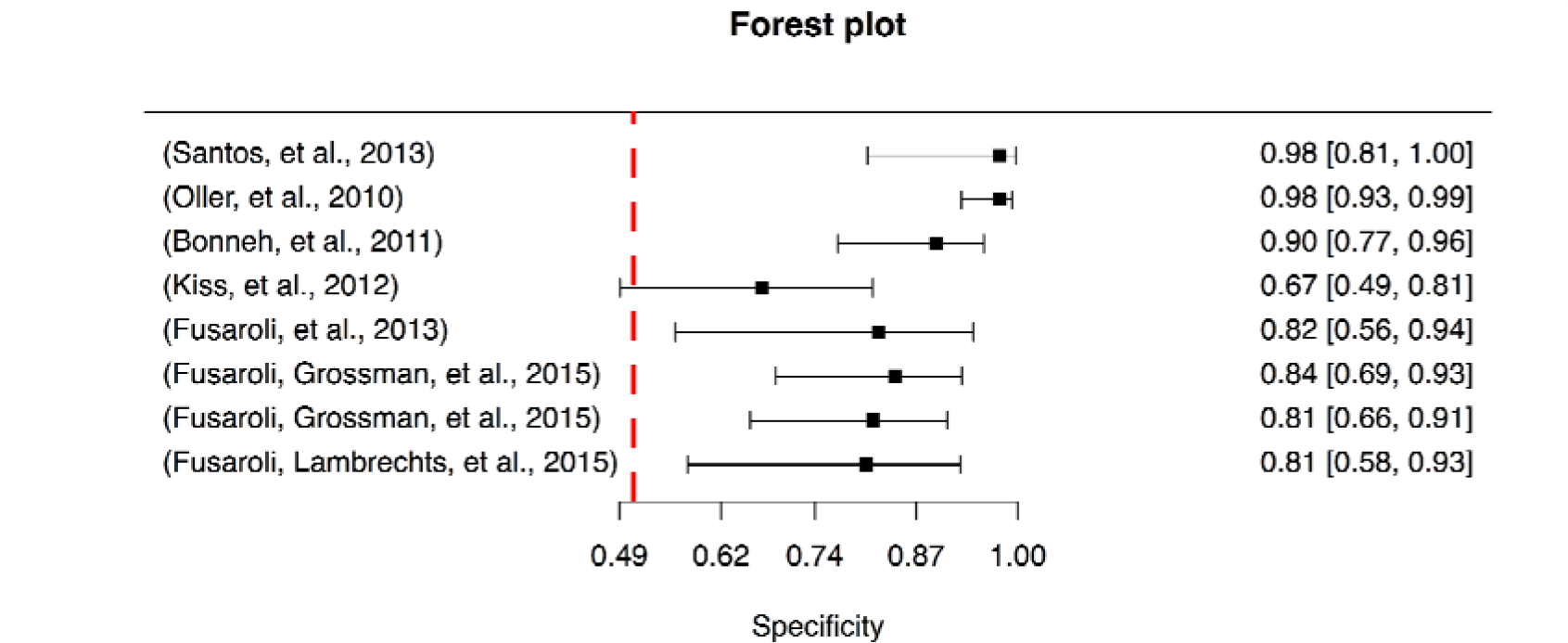
Forest plot of the algorithms' specificities in automatically discriminating between the ASD and comparison populations. The x-axis reports the specificity and the y-axis the studies for which the relevant statistics were available. The dotted line indicates specificity at chance level, that is, 50%.

Besides the classification of voice into ASD and comparison groups, 4 studies demonstrate the possibility of predicting severity of clinical features (ADOS total scores, ADOS Stereotyped Behavior and ADOS Reciprocal Social Interaction) from acoustic measures, in particular pitch, shimmer and jitter (Bone, et al., 2014; Fusaroli, et al., 2013; Fusaroli, Grossman, et al., 2015; Fusaroli, Lambrechts, et al., 2015). However, differences in terms of methods and measures make comparison between studies difficult.

## 6. Discussion

### 6.1 *Overview*

Clinical practitioners have long attributed distinctive voice and prosodic patterns to individuals with ASD (Asperger, 1944; Kanner, 1943). We set out to systematically review the evidence for such patterns and their potential as a marker of ASD. We identified 34 articles involving 30 univariate and 15 multivariate machine-learning studies. Sample sizes were limited, with a mean of 21.14 (SD: 16.36) and a median of 17 (IQR: 10.5) ASD participants across the univariate studies and a mean of 24.1 (SD: 18.24) and a median of 17 (IQR: 15.5) across the multivariate ones.

The univariate studies reported as many null results as significant differences between ASD and comparison groups. Meta-analyses identified reliable, but small effects for pitch mean and range, corresponding to a discriminative accuracy of approximately 61–64%. The multivariate machine-learning studies, by contrast, painted a more promising picture and largely outperformed the univariate ones, with accuracy ranging from 70% to 96% for separating individuals with ASD from comparison participants. The multivariate attempts at predicting severity of clinical features do not systematically outperform the univariate studies (univariate R^2^ between 0.18 and 0.46; multivariate Adjusted R^2^ between 0.13 and 0.8). Whilst the multivariate findings are stronger and involve more robust statistical procedures (such as validation procedures), there has been no general attempt to replicate findings across multiple studies using similar methods. Because of the complexity and heterogeneity of feature extraction, selection and of the statistical models involved in the multivariate studies, it is not possible to assess which (if any) of the acoustic features are most informative for diagnosis and clinical features across studies.

### 6.2. Obstacles in identifying an acoustic marker for ASD

We raised the possibility that acoustic features of vocal production could be used as a marker of ASD. We defined a marker of ASD as a directly measurable index that is derived from sensitive and reliable quantitative procedures and is associated with the disorder and/or its clinical features. We identified as additional challenges the need to assess the heterogeneity of individuals with ASD (e.g. in severity of clinical features) and the progression of clinical features over time (e.g. in presence of intervention program or aging).

We could not identify any single feature that could yet serve the role of a marker. While many aspects of vocal production in ASD have long been described as different, there have been few consistent findings among studies, except for pitch mean and range. The multivariate machine-learning approach to vocal production in ASD seems promising, albeit yet unsystematic; it can capture the complex and often non-linear nature of the acoustic patterns that may give rise to the clinical impression of atypical voice and prosody in ASD. Indeed, such impressions are often based on multiple types of information (Forbes-Riley & Litman, 2004; Liscombe, Venditti, & Hirschberg, 2003).

Many advances have thus been made since McCann & Peppé's (2003) review: a larger number of acoustic features have been quantitatively defined and more complex statistical techniques have been developed. However, the search for a vocal marker of ASD has still to overcome four obstacles: small sample sizes; few replications of effects across studies; too heterogeneous methods for the extraction of acoustic features and their analysis; and limited theoretical background for the research. First, people with ASD present diverse clinical features with different levels of severity. Five of the reviewed studies sought to investigate the relation between severity of clinical features and acoustic patterns. However, because the sample size of each study was too low (median of participants with ASD<30), it is difficult-if not impossible-to control for the large natural heterogeneity among individuals in terms of clinical features and their severity. Second, most of the studies reviewed focused on different acoustic features, which entails that effects rarely are replicated and that it is difficult to perform reliable meta-analyses of effect sizes. Third, the reviewed studies differed considerably with respect to methods and statistical analysis. For example, we identified three types of speech-production task (constrained production, spontaneous production and social interaction), each of which is likely to involve distinct social and cognitive demands and therefore different vocal production patterns, but more fine-grained typologies could be used. This would also enable the assessment of whether acoustic markers of ASD could represent biomarkers, that is, be directly related to underlying biological processes as those involved in respiration and fine-motor control of the vocal folds. Further, different studies not only use different acoustic features but also use different methods for feature extraction - if described at all-making comparisons between studies difficult^6^. This lack of clarity is especially problematic for machine-learning techniques^7^.

A final issue to be mentioned is the relation between acoustic markers, clinical assessment and diagnosis (or clinical features). Would acoustic markers of ASD contribute new information to the clinical assessment? Technically, the machine learning procedures analyzed rely on existing clinical assessment to learn the relation between acoustic features and ASD. In other words, they cannot get better than the clinical assessment they are trained on. Nevertheless, there are several advantages in employing acoustic markers of ASD and its clinical features. First, the identification of acoustic markers would represent a fast, cheap, non-invasive procedure, which could speed up the diagnostic process. Second, the procedure could support the diagnostic process in objective ways, increasing the reliability of the clinical features assessment especially for less experienced practitioners. Third, acoustic markers of ASD and clinical features could point to mechanisms underlying the disorder and its various impairments allowing for a simultaneous assessment of several clinical features and their progression over time. Whether these potentialities can be lived out is still an empirical question, which requires more collaborative and open research processes.

### 6.3. Towards a more collaborative and open research process

The combination of promising results and a lack of a systematic approach is far from rare in the study of acoustic patterns in neuropsychiatric conditions (Cohen, Mitchell, & Elvevåg, 2014; Cummins, et al., 2015; Weed & Fusaroli, submitted). We need to develop a systematic approach to vocal production in ASD, accounting for the heterogeneity of the disorder, the individual differences of the participants and their progression through aging and intervention, for it to be of clinical relevance. To achieve this goal we advocate more open and cumulative research practices. We therefore outline three recommendations for future research: open data, open methods, and theory-driven research.

*Open Data.* Many of the reviewed studies did not report the necessary information for performing meta-analysis. For example, we could not account for the role of age in the patterns observed, as we could not access participant-level data matching acoustic and demographic measures. The field as a whole would benefit from sharing datasets, which would allow for across-study comparisons and for larger scale analyses. While voice recordings are often sensitive data in clinical population, and therefore not easily shareable, the extracted acoustic measures do not always share this restriction. In line with this recommendation, the data used here are available at https://github.com/fusaroli/AcousticPatternsInASD.

*Open Methods.* The quantitative assessment of acoustic measures presents the researcher with several important choices: for example, how should the audio signal be recorded and preprocessed, which parameters should be used to extract the different acoustic features, and whether the extracted data is transformed (e.g. applying a logarithmic transform to fundamental frequency). Recording devices and setup might have a strong impact on the quality of the recording and affect the possibility of extracting source and energy-based measures such as intensity and voice quality (see Orlikoff and Kahane, 1991 and Degottex et al 2014). It is therefore a good practice to ensure that: i) The same device is used for the full data collection and the technical specifics of the device should be reported. ii) The device maintains a constant distance from the speaker's mouth. Recordings from table-top omnidirectional microphones are susceptible to multiple artifacts due to different posture, movements and agitation in the participants with ASD affecting the mean level of sound pressure and its variability. Even when those suggestions cannot be followed (e.g. when an existing clinical corpus is used for the analysis), reporting the recording device and procedure ensures the possibility to choose and assess only appropriate acoustic features, e.g. excluding intensity and voice quality in presence of sub-optimal recordings.

Pre-processing and feature extraction have even more degrees of freedom and detailed reports of the choices are necessary. Otherwise replication and cross-talk between research groups are impossible. Ideally, the full data-processing pipeline should be automated and the script used to do so should be published as supplementary material (or on public code repositories such as GitHub). The literature on vocal production in Parkinson's and affective disorders might serve as an example for researchers investigating vocal production in ASD (Degottex, Kane, Drugman, Raitio, & Scherer, 2014; Tsanas, et al., 2011). In line with this recommendation, the R code employed in this paper is available at https://github.com/fusaroli/AcousticPatternsInASD, and can be easily improved and/or used to update the meta-analysis as new studies are published.

*Theory-driven research.* A common feature of the studies reviewed is the lack of theoretical background. For example, limited attention is paid to clinical features and their severity and the choice of the speech-production task and acoustic measures used is often under-motivated. On the contrary, by putting hypothesized mechanisms to the test, more theory-driven research on vocal production in ASD would improve our understanding of the disorder itself. For examples, recent models of impaired perceptual and motor anticipation in ASD (Palmer, Paton, Kirkovski, Enticott, & Hohwy, 2015; Van de Cruys et al., 2014) would predict the presence of overcorrection in vocal production in ASD (e.g. bursts of jitter and shimmer). Further, models of social impairment in ASD could be tested by analyzing the acoustic dynamics involved in conversations, such as reciprocal prosodic adaptation and compensation (Dale, Fusaroli, Duran, & Richardson, 2013; Fusaroli, Raczaszek-Leonardi, & Tylén, 2014; Fusaroli & Tylén, 2012; Hopkins, Yuill, & Keller, 2015; Lambrechts, Yarrow, Maras, & Gaigg, 2014; Pickering & Garrod, 2004; Slocombe et al., 2013).

In general, different speech-production tasks involve different social and cognitive demands and such differences might account for much of the unexplained variance between the reviewed studies. We therefore recommend data collection using several motivated speech-production tasks, especially combining existing clinical and ecological speech recordings with tasks chosen based on hypothesized mechanisms underlying clinical features. On one hand, structured tasks might allow the researcher to control for confounds and test for the role of specific experimental factors. Further, several standardized tests - including ADOS interviews - involve vocal production and their systematic collection and use could enable the construction of large datasets comparable across labs and languages. On the other hand, structured tasks might not offer representative samples of vocal productions in ASD, as individuals with ASD differ in terms of what they can do if tested and what they actually do in their everyday life (Fine, Bartolucci, Ginsberg, & Szatmari, 1991; Klin, Jones, Schultz, & Volkmar, 2003). Recent technological developments enable unobtrusive longitudinal recordings, opening up for the study of prosody and other social behaviors during everyday life (Vosoughi, Goodwin, Washabaugh, & Roy, 2012; Warlaumont, et al., 2014). This might in turn help us better understand the everyday dynamics of social impairment in ASD.

## 7. Conclusion

We have systematically reviewed the literature on distinctive acoustic patterns in ASD. We did not find conclusive evidence for a single acoustic marker for ASD and predictor for severity of clinical features. Multivariate machine-learning research provides promising results, but more systematic cross-study validations are required. To advance the study of vocal production in ASD, we outlined three recommendations: more open, more cumulative and more theory-driven research.

## Footnotes

1 Additional data were provided by the authors of (Bonneh, Levanon, Dean-Pardo, Lossos, & Adini, 2011; Grossman, et al., 2010), whom we gratefully acknowledge. As this data is fully reported in the publicly accessible dataset, we will not further distinguish it from the data reported in the articles reviewed.

2 HFA indicates High Functioning Individuals with ASD, AS Asperger's Syndrome, PDD-NOS pervasive developmental disorder not otherwise specified. Raven indicates Raven's Coloured Progressive Matrices. PPVT; Clinical Evaluation of Language Fundamentals

3 It should be noted that a few studies attempted to separate different groups within the autism spectrum. One study did not find any significant difference between Asperger Syndrome (AS), high-functioning and pervasive developmental disorder not otherwise specified (PDD-NOS) (Paul, Bianchi, Augustyn, Klin, & Volkmar, 2008). However, another found that individuals with AS produced larger pitch ranges than speakers with PDD-NOS (Kaland, et al., 2012), a pattern repeated when comparing high-with lowerfunctioning people with autism (Depape, et al., 2012).

4 NN: neural networks; SVM: support vector machines; k-NN: nearest neighbors; DA: discriminant analysis. *Accuracy* indicates the percentage of correctly identified data points in the testing set. *Specificity* indicates the ability to correctly identify controls as controls, *Sensitivity* or recall indicates the ability to correctly identify targets as targets. *Precision* indicates the probability that a positive diagnosis does indeed entail the presence of a disorder. For regressions, performance is measured in terms of variance explained, R^2^, which in turn tends to be penalized according to the number of features included, Adjusted R^2^ (Hastie, et al., 2009).

5 Given the heterogeneity of the studies in terms of acoustic measures and algorithms a metaanalysis would not be reliable and is not reported. The curious reader can find the code for performing one at https://github.com/fusaroli/AcousticPatternsInASD

6 For instance, the parameters to define the accepted ceiling of the fundamental frequency might vary from 400 Hz to 700 Hz. Higher ceilings have been shown to better capture acoustic differences features in ASD (Kiss, et al., 2012), however the definition of the ceiling employed is very rarely reported.

7 It has been shown, for example, that recording participants with ASD and comparison participants at different locations (which was unreported) induced artificially high discrimination accuracy due to the properties of each location's background noise (Bone, et al., 2013).

## Acknowledgments

This work was supported by the Seed Funding Program of The Interacting Minds Center, grant “Clinical Voices” (RF) and the Calleva Research Centre for Evolution and Human Sciences (DB).

